# A unified model of amino acid homeostasis in mammalian cells

**DOI:** 10.1101/2021.02.08.430327

**Authors:** Gregory Gauthier-Coles, Jade Vennitti, Zhiduo Zhang, William C. Comb, Kiran Javed, Angelika Broer, Stefan Broer

## Abstract

Homeostasis is one of the fundamental concepts in physiology. Despite remarkable progress in our molecular understanding of amino acid transport, metabolism and signalling, it remains unclear by what mechanisms cytosolic amino acid concentrations are maintained. We propose that amino acid transporters are the primary determinants of intracellular amino acid levels. We show that a cell’s endowment with amino acid transporters can be deconvoluted by a logical series of experiments. This was used to computationally simulate amino acid translocation across the plasma membrane. For two different cancer cell lines and human myotubes, transport simulation generates cytosolic amino acid concentrations that are close to those observed *in vitro*. Perturbations of the system were replicated *in silico* and could be applied to systems where only transcriptomic data are available. The methodology developed in this study is widely applicable to other transport processes and explain amino acid homeostasis at the systems-level.

## Introduction

The concept of homeostasis is a cornerstone of physiological research. Similar to other metabolites, plasma and cellular amino acid concentrations are kept within their physiological range. After a meal, amino acid concentrations rise, but return to homeostatic levels within hours ^1,2^. In contrast to glucose, which is found at lower than plasma concentrations in the cytosol, due to rapid conversion into glucose-6-phosphate ^3^, cytosolic amino acid levels are several-fold higher than those in blood plasma ^4^. This provides sufficient amino acid pools to support protein biosynthesis and contributes to the osmotic pressure of the cytosol ^5^.

Two signalling systems regulate amino acid homeostasis, namely mTORC1 and GCN2/ATF4. When cytosolic and lysosomal amino acid levels are sufficient, they activate the mTORC1 complex ^6^, needed for cell division, efficient translation and ribosome production ^7^. Depletion of amino acids switches off mTORC1, but also generates a more specific response to deal with amino acid imbalances via the GCN2/ATF4 pathway of the integrated stress response ^8^.

Amino acid transport in mammalian cells is mediated by more than 60 different uniporters, symporters and antiporters ^4,9^. Uniporters for amino acids do not occur frequently in mammalian cells, because they equilibrate cytosolic and plasma amino acid concentrations, thus preventing elevated levels of amino acids in the cytosol. Exceptions are cationic amino acid transporters, such as CAT1 (SLC7A1) ^10^, which accumulate their substrates due to a positive charge. Examples of widely expressed symporters are the Na^+^-neutral AA cotransporters SNAT1 (SLC38A1) and SNAT2 (SLC38A2) ^5,11^. The most numerous group of amino acid transporters are antiporters ^12^. Examples are LAT1 (SLC7A5, large neutral amino acids) y^+^LAT2 (SLC7A6, neutral and cationic amino acids), ASCT1 (SLC1A4, small neutral) and ASCT2 (SLC1A5, polar neutral)^13^. Although antiporters cannot mediate net uptake, they are upregulated in many fast growing cell types, especially cancer cells ^13^. Cationic amino acids can enter cells through uniporters such as CAT1 (SLC7A1), or through antiporters, such as y^+^LAT2 (SLC7A6), which exchanges a neutral amino acid together with Na^+^ or K^+^ against a cationic amino acid ^14^. Cysteine is acquired by cells as a neutral amino acid through symporters, but can also be acquired as negatively charged cystine (Cys-S-S-Cys) in exchange for glutamate through the antiporter xCT (SLC7A11)^15^. Some of these transporters are actively regulated by ATF4/GCN2 and mTORC1, such as CAT1 (SLC7A1), xCT (SLC7A11), and SNAT2 (SLC38A2) ^16^.

A complete analysis of amino acid transport activities in a given cell type has been difficult to achieve due to overlapping substrate specificities, different affinities and unclear subcellular localisation. It is also unclear how these transporters work together to provide cells with a harmonized mix of amino acids to sustain protein translation and other critical amino acid dependent functions.

In a recent review Wolfson and Sabatini ^17^ state “it is not known how concentrations of amino acids in the media correlate with intracellular, specifically cytosolic, amino acid concentrations”. Here we show that this correlation is determined by a combination of transport processes, which generate a stable equilibrium. The amino acid transporter combination of a cell can be deconvoluted using flux experiments and stable isotope labelling to derive kinetic parameters. These data can then be used as inputs for in silico simulation of amino acid transport, thereby providing a systems level understanding of amino acid homeostasis.

## Results

### Amino acid transporter composition in two cell line

We chose two human cancer cell lines, namely A549 lung cancer cells and U87-MG glioma cells as models for this study, because they are widely used and have different tissue origins. To qualitatively survey the amino acid transporter set in each cell line, we analysed public databases and used RT-PCR to determine the expression of amino acid transporters at the transcriptomic level (Fig. 1 and Fig. S1). For each transporter an expression score was generated. In the second step, we used antibodies combined with surface biotinylation to identify transporters that are potentially active (Fig.1 and Fig. S2/S3). Surface biotinylation/western blotting is an important verification for two reasons. Firstly, mRNA levels did not always correlate with surface expression. For instance, SNAT2 (Na^+^-AA symporter for polar neutral AA) mRNA is abundant, but its protein can barely be detected at the cell surface as shown previously in 143B cells ^18^. This was also observed for A549 and U87-MG cells (Fig.1 and Fig. S2/S3). SNAT5 (SLC38A5, Na^+^-AA symporter/H^+^ antiporter for polar neutral AA), by contrast, has a strong signal in western blots, but its mRNA was barely detectable. Secondly, some transporters are predominantly lysosomal, such as PAT1 (SLC36A1, Proton AA symporter for small neutral AA), but may also occur at the cell surface in some cell types ^19^. Transporters strongly expressed at the cell surface in both cell lines were: ASCT2 (antiporter for polar neutral AA), 4F2hc (SLC3A2, transporter-associated trafficking subunit) and its associated transporters LAT1 (antiporter for large neutral AA), y^+^LAT2 (antiporter for neutral (+Na^+^) and cationic AA), xCT (glutamate-cystine antiporter), SNAT1 (Na^+^-AA symporter for polar neutral AA) and SNAT5. In addition, there were cell-specific transporters such as B^0^AT2 (SLC6A15, Na^+^-AA symporter for neutral AA) and ASCT1 (antiporter for small polar neutral AA) in U87-MG cells. Dedicated anionic amino acid transporters are not essential as aspartate and glutamate can be generated metabolically. The initial survey simplifies the design and interpretation of the radioactive flux experiments outlined in the next paragraph.

**Fig. 1.**
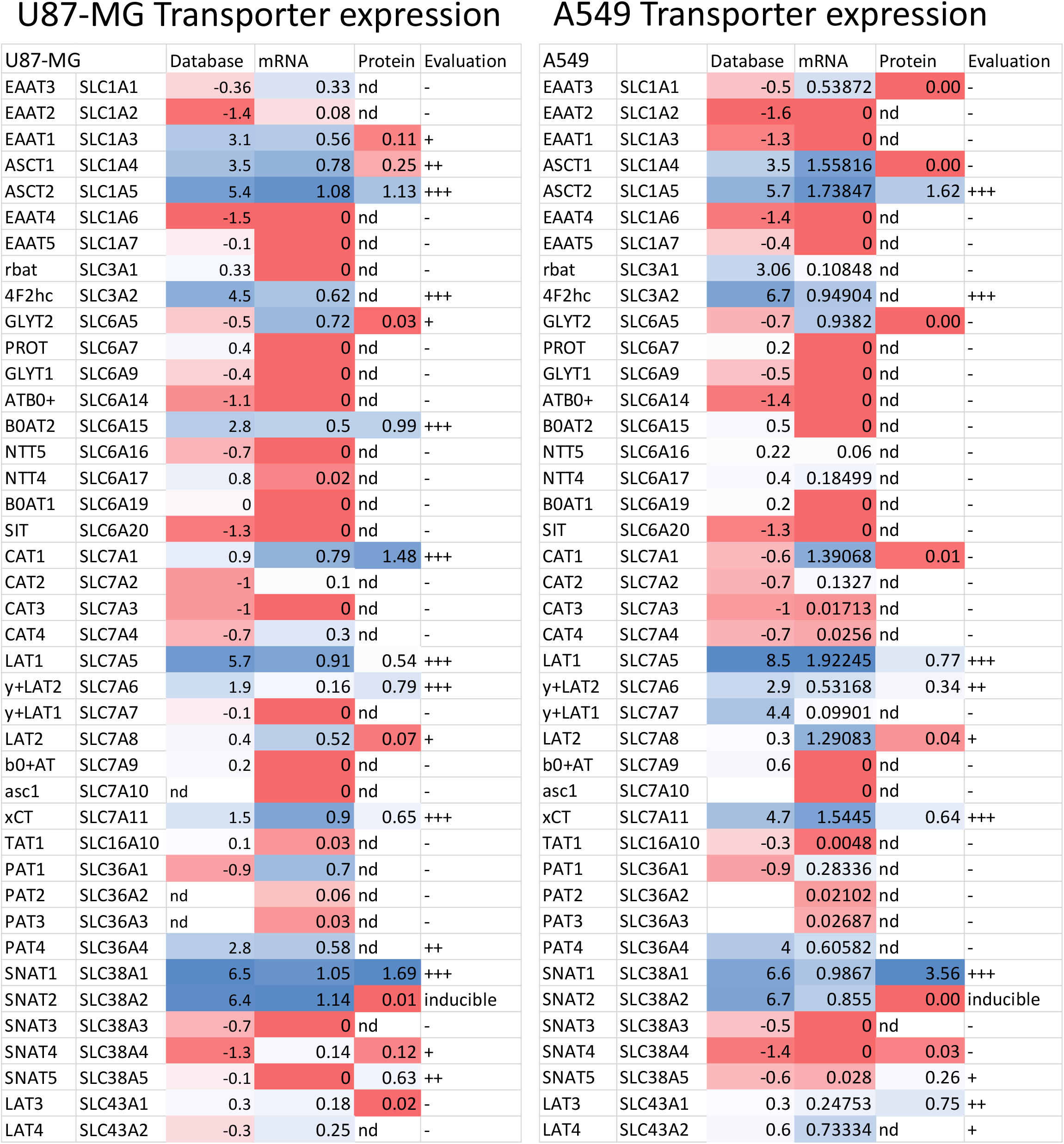
Evaluation of transporter expression in U87-MG glioma and A549 lung carcinoma cells. The mRNA expression levels were compiled from Oncomine (log2 normalised mRNA expression) and also analysed by RT-PCR (normalised to clathrin intensity). Blue indicates high level of expression, red indicates low levels of expression. High levels of mRNA expression were correlated to western blotting (normalised to Na^+^/K^+^-ATPase intensity). Overall evaluation was based on the combined dataset. +++ high activity expected, ++ activity expected to be readily detectable, + marginal activity, -no activity expected.

### Amino acid transport activity in U87-MG cells

Quantification of amino acid transporter expression in this study was derived from transport experiments using radiolabelled amino acids. Table S1 lists all transport conditions to identify specific transporters. Amino acid transporters typically accept groups of related amino acids, such as large non-polar neutral, small neutral, cationic, and anionic ^20^. To capture all amino acid transporters in a given cell, we used substrates to cover these groups, namely leucine, glutamine, alanine, arginine, glycine, proline and glutamate. To discriminate between individual transporters, ion dependence, inhibitors and amino acid competition assays were used. A breakdown of leucine transport in U87-MG cells is shown in Fig. 2A. Notably, leucine transport was largely mediated by the Na^+^-independent transporter LAT1, as evidenced through its inhibition by the specific inhibitor JPH203 ^21^. The Na^+^-dependent uptake of leucine in U87-MG cells was mediated by the branched-chain amino acid transporter B^0^AT2 and by the amino acid exchanger y^+^LAT2 -the only Na^+^-dependent BCAA transporters expressed in U87-MG cells. Their activity can be deduced from the fraction inhibited by loratadine (B^0^AT2) ^22^ and arginine (y^+^LAT2). Accordingly, leucine uptake was completely abolished in the presence of JPH203 in Na^+^-free transport buffer. SNAT2 can also mediate leucine transport, but its contribution as assessed by the amino acid analogue *N*-Methyl-aminoisobutyric acid (MeAIB) ^23^ was too small to be significant. MeAIB is a well-known inhibitor of SNAT1/2/4 (SNAT4, SLC38A4: Na^+^-AA symporter for polar neutral AA). The non-specific amino acid analogue 2-amino-2-norbornane-carboxylic acid (BCH) inhibits LAT1/2 and B^0^AT2, thereby explaining its strong effect on leucine uptake. In combination, this accounted for all of leucine uptake. We did not analyse aromatic amino acid transport separately, because of the complete overlap with leucine accepting transporters in U87-MG cells.

**Fig. 2.**
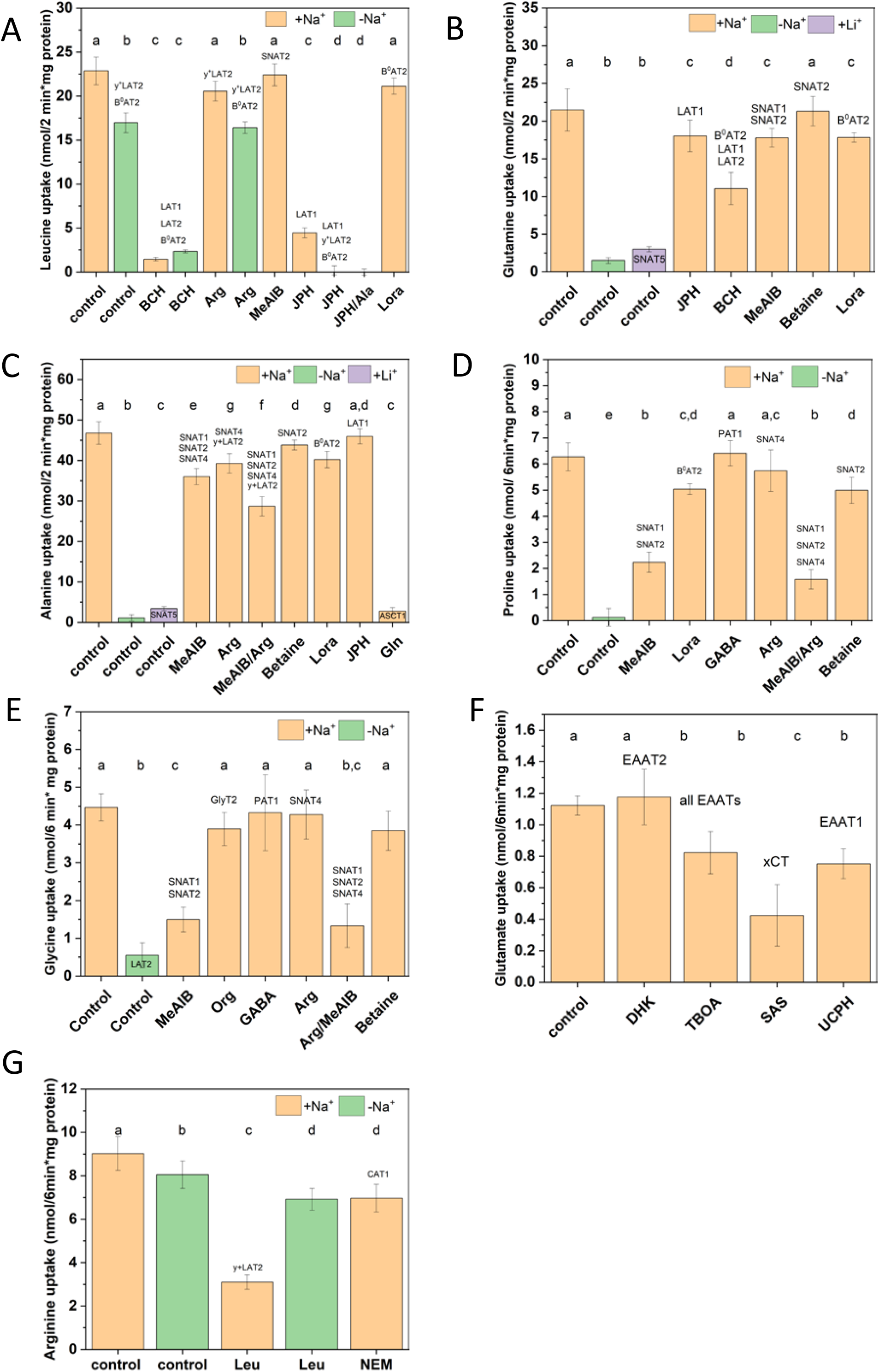
Deconvolution of amino acid transport activities in U87-MG glioma cells. The transport of representative [^14^C]amino acids (A) leucine (100 µM, 2 min, n=15), (B) glutamine (100 µM, 2 min, n=12), (C) alanine (300 µM, 2 min, n=9), (D) proline (100 µM, 6 min, n=21), (E) glycine (100 µM, 6 min, n=24), (F) glutamate (100 µM, 6 min, n=9) and (G) arginine (100 µM, 6 min, n=9) is shown. AA uptake was measured in Na^+^-based buffer (orange), NMDG^+^-based buffer (green) or Li^+^-based buffer (purple). In the control experiments no inhibitor or competitor was added. With the exception of NEM (10 min preincubation), inhibitors and competitors were added together with the substrate at the concentrations given in Table S3. To facilitate analysis, blocked transporters relevant to the cell line are shown above the bars, remaining transporters are shown in the bar where indicated. Non-specific uptake/binding was evaluated by addition of 10 mM unlabelled substrate and was subtracted from all experiments. ANOVA with Tukey’s multiple comparison test was used to analyse differences between groups. Groups that were not significantly different from each other at p<0.05 were assigned the same letter. Abbreviations: Amino acids are shown in three-letter code; BCH, 2-amino-2-norbornane-carboxylic acid; DHK, dihydrokainate, JPH, JPH203; Lora, loratadine; MeAIB, *N*-methyl-aminoisobutyric acid; NEM, *N*-ethylmaleimide; Org, ORG25543; SAS, Sulfasalazine, TBOA, DL-threo-β-benzyloxyaspartate; UCPH, UCPH101.

In contrast to leucine transport, uptake of glutamine was largely mediated by Na^+^-dependent transporters (Fig. 2B). Glutamine is a poor substrate of LAT1 and accordingly, inhibition by JPH203 was small. BCH, which also inhibits LAT2 and B^0^AT2, showed a stronger effect than JPH203. A fraction of glutamine uptake was inhibited by MeAIB, indicating it was facilitated by either SNAT1 or SNAT2. Betaine, which discriminates between SNAT1 and SNAT2, had no effect, suggesting SNAT1 to be the dominant MeAIB-sensitive glutamine transporter. This is also in agreement with the lack of MeAIB inhibition on leucine transport. Loratadine was used to identify the fraction of glutamine uptake mediated by B^0^AT2 ^22^. A particular feature of glutamine transporters SNAT3 (SLC38A3) and SNAT5 (SLC38A5) is their ability to retain transport activity when NaCl is replaced by LiCl ^11^. This was supported by glutamine transport in LiCl (purple bar) being higher than the glutamine transport in the absence of Na^+^ (green bar). In the absence of SNAT3 expression, this component was attributed to SNAT5. All of glutamine transport was sensitive to inhibition by alanine, suggesting that the remaining Na^+^-dependent uptake was mediated by ASCT2, a dominant neutral amino acid transporter in all cancer cells ^13^. Despite significant efforts, reliable specific inhibitors of ASCT2 are yet to be developed ^24^. As a result, the remaining activity was allocated to ASCT2. These experiments account for all of glutamine uptake, and with the exception of SNAT5, concur with mRNA expression data.

Alanine uptake (Fig. 2C) was also largely Na^+^-dependent and mediated by the same range of transporters as glutamine uptake. It was partially sensitive to inhibition by MeAIB (SNAT1/2/4), arginine (SNAT4 and y^+^LAT2), and loratadine (B^0^AT2), but not by JPH203 (LAT1) consistent with the substrate specificity of LAT1. Thus, the small uptake activity observed in the absence of Na^+^ was assigned to LAT2. The related transporter ASCT1 excludes glutamine and as a result, we assigned the glutamine-resistant portion of alanine transport to ASCT1. A small lithium-dependent component was attributed to SNAT5. The large remaining fraction was attributed to ASCT2. These experiments accounted for all of alanine uptake.

Proline (Fig. 2D) and glycine (Fig. 2E) often have specific transporters, because they tend to be poor substrates of other amino acid transporters. Proline transport was entirely Na^+^-dependent, excluding proton-dependent transporters of the SLC36 family (PAT1-4, SLC36A1-4). This simplifies the interpretation of MeAIB inhibition, which affects SNAT1/2/4 and PAT1/2. Inhibition by betaine was less strong than by MeAIB suggesting that SNAT1 was the dominant proline transporter. We attributed the arginine-sensitive portion of proline uptake to SNAT4. Moreover, the combination of MeAIB and arginine is more powerful than MeAIB alone. GABA did not inhibit proline transport, confirming the absence of PAT1. The fraction of proline transport inhibited by loratadine was attributed to B^0^AT2. Together these transporters explained proline uptake in U87-MG cells. Because loratadine has not been extensively tested against other transporters, we confirmed the contribution of B^0^AT2 to proline, alanine and glutamine transport by RNAi mediated silencing, which matched the effect of loratadine (Fig. S4). Glycine transport was also dominated by SNAT1 and SNAT2, but glycine showed residual transport in the absence of Na^+^, which was assigned to LAT2. Inhibition by arginine was not significant suggesting that SNAT4 did not contribute to glycine uptake. A small fraction of glycine transport was inhibited by ORG24453 ^25^, which was consistent with a weak protein expression of GlyT2 (Fig. S2).

Glutamate uptake (Fig. 2F) was mediated by xCT, which is sensitive to inhibition by sulfasalazine ^26^, and to a smaller extent by EAAT1, which is sensitive to inhibition by UCPH101 ^27^. The inhibition by UCPH matched the fraction of transport blocked by the non-specific excitatory amino acid transporter (EAAT) inhibitor DL-*threo*-β-benzyloxyaspartate (TBOA) ^28^, suggesting that no other EAAT was involved.

Arginine uptake (Fig. 2G) was entirely Na^+^-independent. A significant fraction of arginine transport was inhibited by leucine in the presence of Na^+^, but not in its absence, a hallmark of y^+^LAT2. This transporter is resistant to treatment with *N*-ethylmaleimide (NEM), while the canonical cationic amino acid transporters CAT1/2 are sensitive. Due to the relative abundance of its mRNA, the NEM-sensitive portion was assigned to CAT1. The combination of these transporters fully accounted for arginine uptake.

### Amino acid transporter activity in A549 cells

Expression of mRNA and protein suggested a slightly simpler array of amino acid transporters in A549 cells, including ASCT2, LAT1/2/3/4, y^+^LAT2, CAT1, xCT, SNAT1/2/5 (Fig. 1). We used the same strategy to analyse amino acid uptake in the human lung cancer cell line A549 (Fig. 3). Leucine transport (Fig. 3A) was mediated by LAT1, LAT2/3/4, y^+^LAT2 and SNAT2. LAT1, LAT2, ASCT2, SNAT1/5 were the main glutamine transporters (Fig. 3B) and alanine transporters (Fig. 3C). Proline uptake was mediated by SNAT1/2/4 (Fig. 3D), while glycine uptake was mediated by SNAT1/2, and ASCT2 (Fig. 4E). Glutamate transport was carried entirely by xCT (Fig. 3F). Arginine uptake was mediated by y^+^LAT2 and CAT1 (Fig. 3G).

**Fig. 3.**
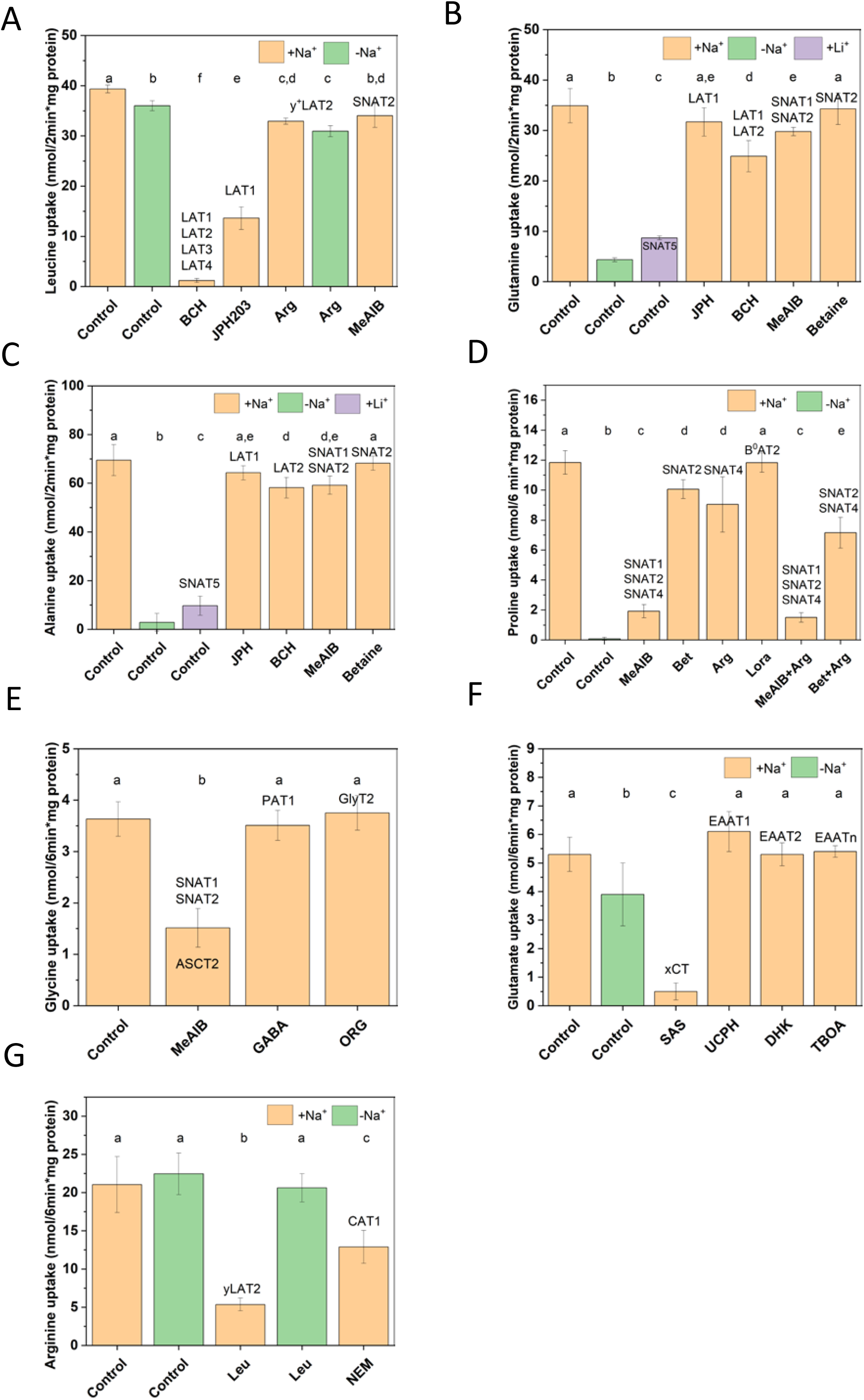
Deconvolution of amino acid transport activities in A549 lung carcinoma cells. The transport of representative [^14^C]amino acids (A) leucine (100 µM, 2 min, n=9), (B) glutamine (100 µM, 2 min, n=9), (C) alanine (300 µM, 2 min, n=9), (D) proline (100 µM, 6 min, n=9), (E) glycine (100 µM, 6 min, n=9), (F) glutamate (100 µM, 6 min, n=9) and (G) arginine (100 µM, 6 min, n=9) is shown. AA uptake was measured in Na^+^-based buffer (orange), NMDG^+^-base buffer (green) or Li^+^-based buffer (purple). In the control experiments no inhibitor or competitor was added. Except for NEM (10 min preincubation), inhibitors and competitors were added together with the substrate at the concentrations given in Table S3. To facilitate the analysis, blocked transporters relevant to the cell line are shown above the bars, remaining transporters are shown in the bar where indicated. Non-specific uptake/binding was evaluated by addition of 10 mM unlabelled substrate and was subtracted from all experiments. ANOVA with Tukey’s multiple comparison test was used to analyse differences between groups. Groups that were not significantly different from each other at p<0.05 were assigned the same letter. Abbreviations: Amino acids are shown in three-letter code; BCH, 2-amino-2-norbornane-carboxylic acid; DHK, dihydrokainate, JPH, JPH203; Lora, loratadine; MeAIB, *N*-methyl-aminoisobutyric acid; NEM, *N*-ethylmaleimide; Org, ORG25543; SAS, Sulfasalazine, TBOA, DL-threo-β-benzyloxyaspartate; UCPH, UCPH101.

**Fig. 4.**
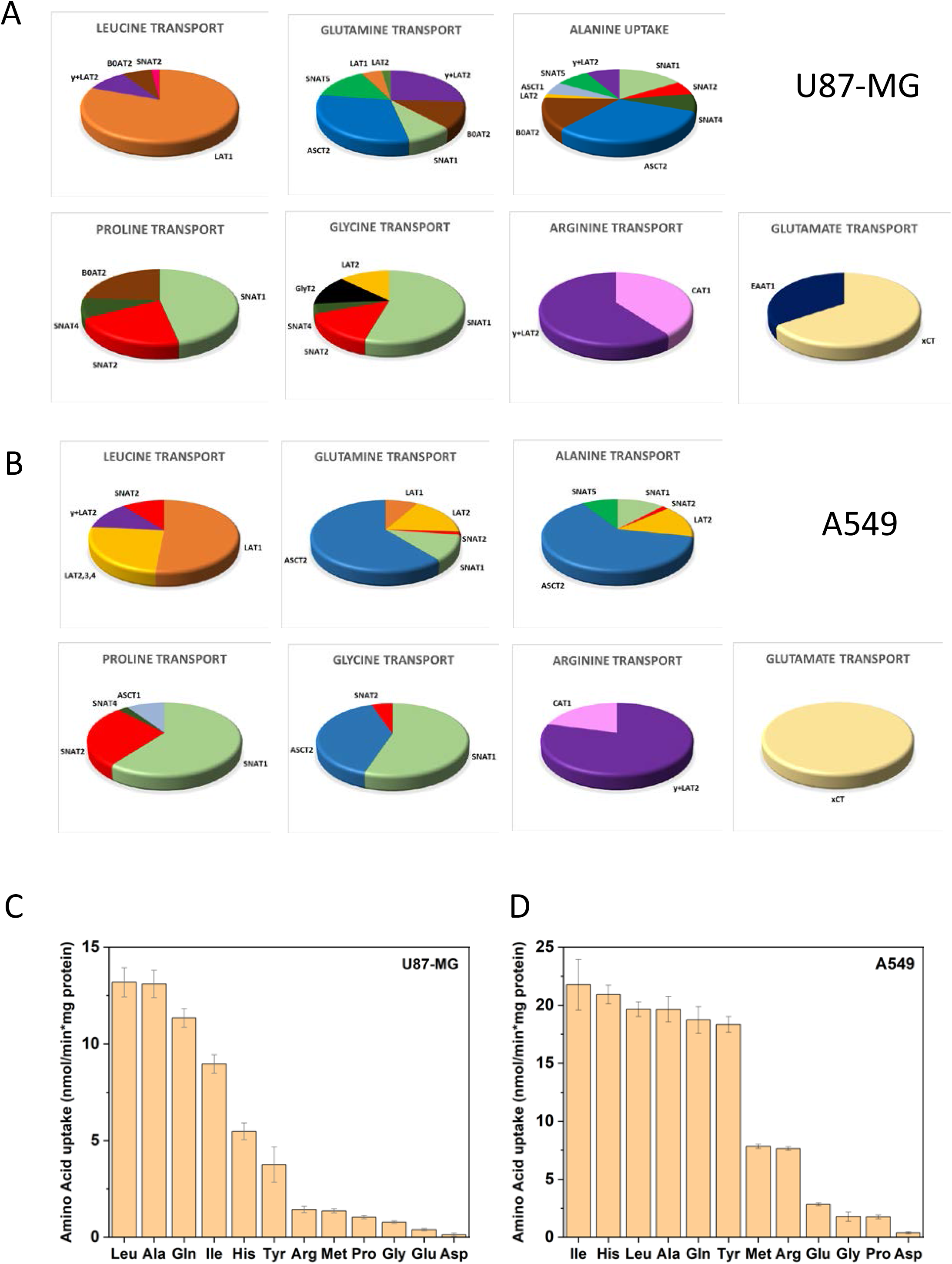
Quantification of plasma membrane amino acid transport. Data from Fig. 3 and Fig. 4 were converted to pie charts, representing the contribution of individual transporters to the uptake of the indicated amino acid. Panel A shows AA transport activities in U87-MG cells, panel B in A549 cells. To compare the uptake activity for a variety of amino acids, uptake was measured at 100 µM in U87-MG cells (C, n=9) and A549 cells (D, n=9).

### Overview of transporter activity

Figure 4 shows a compilation of all active amino acid transporters and the allocation of fluxes in U87-MG cells (Fig. 4A) and in A549 cells (Fig. 4B). The data confirm the dominance of the antiporters LAT1 for leucine, ASCT2 for glutamine and alanine, and xCT for glutamate. Notably, uptake via xCT is non-productive glutamate/glutamate exchange, because cystine is reduced to cysteine in the cytosol, leaving only glutamate as an exchange substrate.

In both cell lines, transport of glutamine, alanine and leucine was much faster than that of arginine, proline and glutamate when measured at a concentration of 100 μM (Fig. 4C,D). Aspartate transport was barely detectable, indicating very low levels of EAATs. This dataset provided the basis of a systems level simulation of cellular amino acid transport to understand their role in amino acid homeostasis.

### Computational simulation of cellular amino acid transport

Amino acid transport is governed by Michaelis-Menten kinetics and electrochemical driving forces. As a result, transporter simulations need to follow kinetic and thermodynamic principles. The transport mechanism for each known amino acid transporter is shown in Table S2. A curated list of K_M_-values of human amino acid transporters is shown in Table S3. These K_M_-values and the V_max_-values derived from the transport analysis were used to simulate amino acid transport in mammalian cells using a number of established principles:

1. Saturation of the transporter by each of its substrate amino acids (AA_i_) follows a simple binding algorithm:

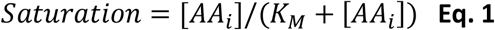
2. Competition between substrates is incorporated by calculation of an apparent K_M_:

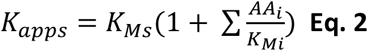
3. A fractional saturation is calculated for each amino acid (AA_i_) using its apparent K_M_.
4. Saturation of the transporter by cotransported ions follows the Hill-equation:

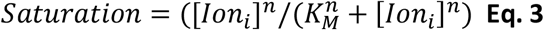
5. Translocation of a charged complex is affected by the membrane potential ^29^. The transport rate was multiplied by 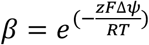 (**Eq.4**) when translocation is favoured by the membrane potential and divided by β, when translocation occurs against the membrane potential.

Using these principles, we devised a set of equations for any kind of transport process (Table S4). A couple of restrictions apply to avoid violating the 2^nd^ law of thermodynamics.

1. Uniporter: The extracellular and intracellular K_M_-values must be equal to avoid accumulation without a substrate gradient.
2. Symporter: The extracellular and intracellular K_M_-values must be equal to avoid accumulation of substrate when the membrane potential is 0 and no substrate gradient is applied. Moreover, β (and 1/β) must decrease as the substrate concentration rises on the opposite side, because the transporter will increasingly operate in electroneutral exchange mode. This is achieved by multiplication of β (or 1/ β) with the fraction of transporters that can carry out net transport. Any symporter must not exceed thermodynamic limits of accumulation.
3. Antiporter: The overall exchange is dependent on the occupancy of the transporter on both sides of the membrane. No transport may occur in the absence of substrate on either side.
4. All transporters: The same calculated V_max_ is applied to the forward and backward flux.

With these rules applied, we devised a computational model, which was implemented in MATLAB (called JDFC from hereon) and uses the following inputs:

1. K_M_-values for each substrate of each expressed transporter (Table S3). Missing K_M_-values were estimated by interpolation.
2. Calculated V_max_ values derived from the experiments outlined above (Table S5).
3. Extracellular amino acid concentrations were fixed according to the media formulation.
4. Initial intracellular amino acid concentrations are variables but were based on actual measurements. The equilibrium is independent of the initial concentrations.
5. Cell volume as determined by Coulter counter analysis (Fig. S5). No correction for organellar space was made, which is <10% v/v in A549 cells ^30^.
6. Optional: Fractional conversion of one amino acid into another (e.g. glutamine into glutamate).
7. Optional: Fractional depletion of an amino acid (metabolism or protein synthesis).
8. A time step used for looping the algorithm.
9. The total number of iterations.

JDFC employs a set of functions for each type of transporter (Table S2/S4). After compiling all functions for a set of transporters in a given cell line, JDFC calculates the fractional saturation of each substrate for each transporter. The corresponding portion of the total flux is allocated to each amino acid. For each substrate amino acid, a small flux was calculated for each participating transporter per time step. These values are then divided by the cell volume to generate incremental changes of AA concentrations, which are summed up for all amino acids. The workflow of the program is shown in Fig. 5A. We hypothesized that this system is inherently stable and comes to an equilibrium point, a condition for a homeostatic process. This was indeed the case (Fig. S6A). Two exceptions were glutamate and aspartate in the presence of an active EAAT. These transporters can theoretically accumulate up to 100M of intracellular glutamate at plasma concentrations. To avoid osmotic stress, both cell lines expressed only minimal levels of EAATs. Instead, glutamate was generated by metabolic conversion of glutamine, which we quantified using [^13^C]glutamine. In U87-MG cells, intracellular glutamine was 90% labelled after 30 min and more than 95% labelled within an hour (Fig. 5B), consistent with fast transport and exchange. Cytosolic glutamate was 30% labelled within 30 min, increasing to 55% within 2h (Fig. 5B). Thus, significant production of glutamate occurs metabolically. In agreement with a functional glutaminolysis pathway, glutamate was metabolised through part of the TCA cycle to form aspartate. Of the initial glutamine, 20% was converted into aspartate in 24h. The maintenance of aspartate and glutamate levels by metabolic conversion was further illustrated by using CB-839 ^31^ to inhibit glutaminase in A549 cells (Fig. 5C). This caused glutamate levels to drop by 53% and aspartate to drop by 60% within 1h, while glutamine increased by 46%. These mechanisms prevent excessive accumulation of aspartate and glutamate through accumulative transport. Moreover, xCT provides a significant pathway for glutamate efflux in exchange for cystine.

**Fig. 5.**
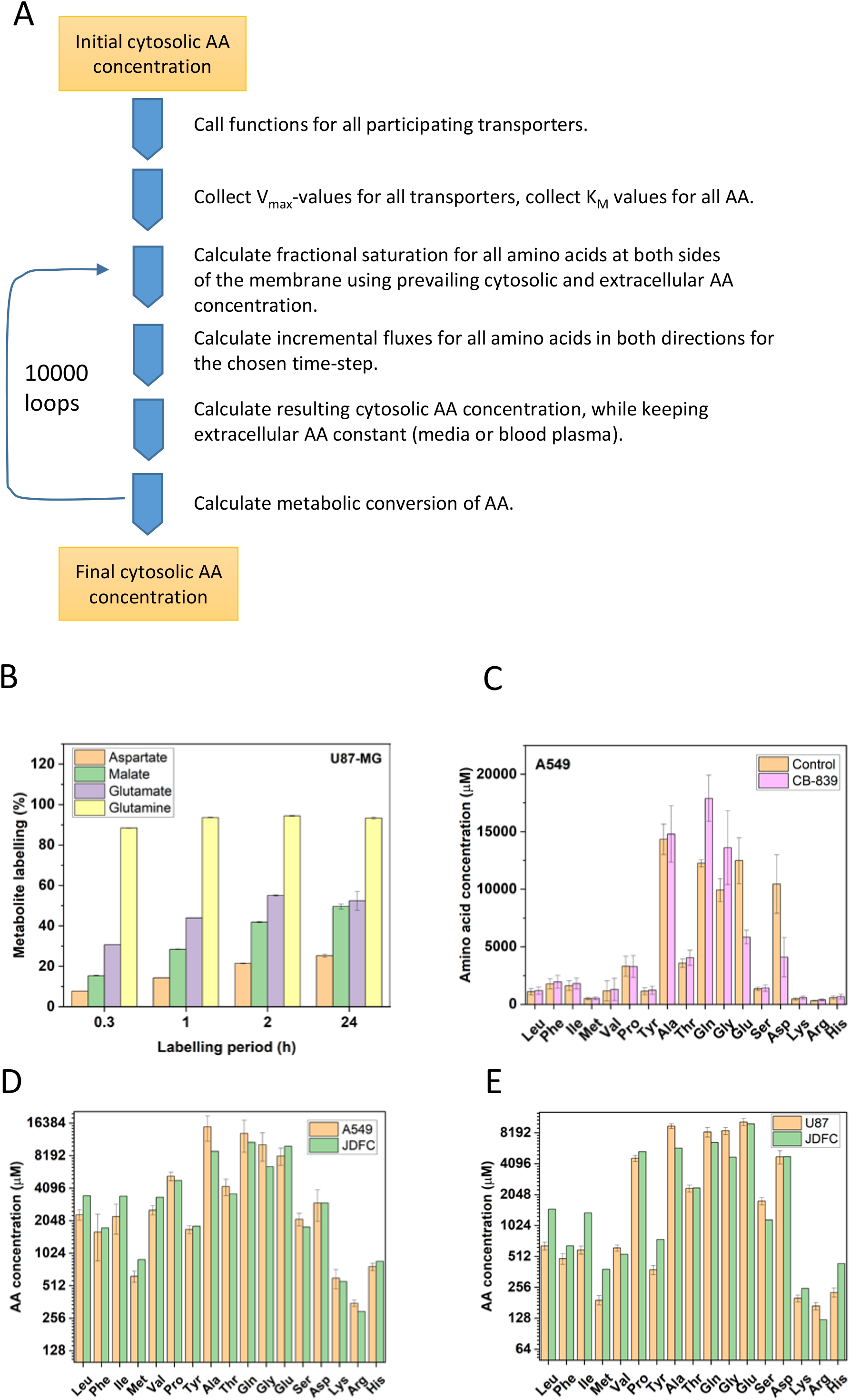
Computational simulation of amino acid transport and metabolomics. A) Principal steps for each iteration of the transport simulation process. For details see text. B) Labelling of metabolic intermediates after incubation of U87-MG cells with [^13^C]glutamine (1 mM, 100% labelled). Enrichment of [^13^C]metabolites is shown after 20, 60, 120 min and 24h. (C) Cytosolic amino acid concentrations were determined by LC-MS. Glutaminase was inhibited in A549 cells by incubation with CB-839 (10 μM, 1h) and amino acid levels compared to the control. (D, E) Cytosolic amino acid concentrations were determined by LC-MS and compared to simulated (JDFC) amino acid concentrations in A549 cells (D) (n=9) and U87-MG cells (E) (n=9). Cells were incubated in BME medium with non-essential amino acids for 30 min before analysis.

### Comparison of *in silico* with *in vitro* data

In view of the stable equilibrium reached by the simulation, we wondered whether this would be accurately reflected in vitro. Thus, we compared simulated and experimental intracellular amino acid concentrations in A549 (Fig. 5D) and U87-MG cells (Fig. 5E). Both datasets were in good agreement with experimental values as assessed by a correlation plot between experimental and predicted values, giving a Pearson’s correlation coefficient of 0.9996 for A549 cells and 0.9588 for U87-MG cells (Fig. S6B/C) over a concentration range from 100-16000 μM.

The endowment of cells with a mix of symporters, antiporters and uniporters links different amino acid pools. We thus tested how amino acid concentrations would react to a fivefold spike of one particular amino acid. To this end, we equilibrated U87-MG cells in a medium containing all amino acids at a concentration of 100 μM and spiked specific amino acids to 500 μM. Interestingly, the spiking affected only the selected amino acid (Fig. 6A), but not any other amino acids. The experiment was fully replicated by the simulation (Fig. 6B), giving a Pearson’s correlation coefficient of 0.968. Consistent with glutamate levels being determined by metabolism, they remained largely unchanged, except when glutamine was spiked. Our model predicts that SNAT1/2/4 accumulate amino acids in the cytosol and the imported amino acids are then used to bring in other amino acids through exchange processes, i.e. tertiary active transport (Fig. 6C). Consistently, inhibition of SNAT1/2/4 by MeAIB resulted in a strong reduction of their substrates glutamine, threonine, methionine and serine, and smaller reductions of alanine, glycine and histidine in A549 cells (Fig. 6D). Glutamate was metabolically generated from glutamine and likewise reduced in the presence of MeAIB. Moreover, through tertiary transport, LAT1 substrates isoleucine, leucine, valine, phenylalanine and tyrosine were reduced, as well. The increase of glutamine in the presence of JPH203 suggests that glutamine is the main efflux substrate in A549 cells. Inhibition of LAT1 by JPH203 (Fig. 6D) resulted in a more selective reduction of valine, tyrosine, phenylalanine, leucine, isoleucine, and histidine. A similar pattern was observed in U87-MG cells (Fig. S6D).

**Fig. 6.**
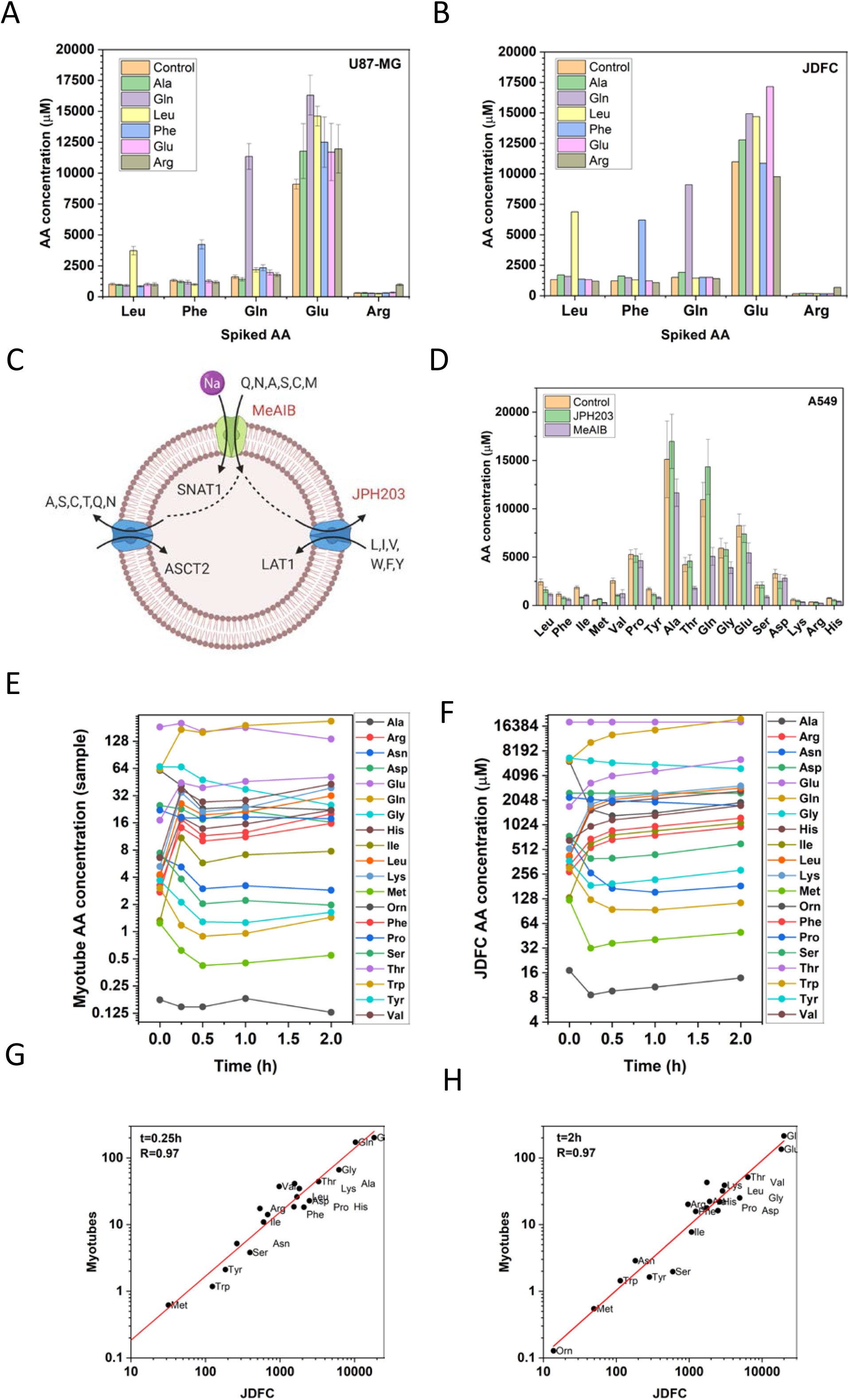
Comparison of in silico and experimental data. U87-MG cells were incubated in a mix of all proteinogenic amino acids (100 µM). The indicated amino acid was spiked at a concentration of 500 µM. Cytosolic amino acid concentrations were determined by LC-MS in U87-MG cells (A) and simulated using JDFC (B). Cells were incubated for 30 min in BME plus non-essential amino acids in the absence (control) and the presence of LAT1 inhibitor JPH203 and the SNAT1/2 inhibitor MeAIB (Scheme in panel C, experimental values in panel D). Amino acids were measured by LC-MS after 30 min. (E) Human primary myotubes were treated with an amino acid mixture corresponding to the constituent amino acids of AXA 2678 for 2h. Cytosolic amino acid concentrations were determined by LC-MS before addition of amino acid mixtures and for 2h after addition. (F) The same initial concentrations were used for JDFC and the simulation was run for 3200 cycles. (G,H) Correlation of experimental and simulated data after t=0.25h, R=0.97 (G) and t=2h, R=0.97 (H).

Amino acid mixtures are being developed as therapeutic products to rebalance metabolic dysfunction such as muscle atrophy ^32^. It is often difficult to predict cellular amino acid changes after consumption of amino acid mixtures. A complete analysis requires transport activity data, while mRNA expression data are readily available for many cell types and tissues. We thus investigated whether relative expression data were sufficient to model amino acid homeostasis in a medically relevant cell model. In this experiment, we used primary human myotubes that were incubated with an amino acid mixture containing leucine, isoleucine, valine, arginine, glutamine, histidine, lysine, phenylalanine, threonine, and *N*-acetyl-cysteine corresponding to AXA2678 (Axcella Health Inc) for the improvement of muscle function ^32^. Cytosolic amino acid concentrations were measured before addition of the amino acid mixture and during a time-course of 120 min (Fig. 6E). Due to a lack of volume data on differentiated myotubes, sample amino acid concentrations are shown. Figure 6F shows the corresponding simulation (excluding N-acetyl-cysteine) involving 25 different amino acid transporters expressed in these cultures. For the modelling, we used transport activities proportional to mRNA expression. The predicted cytosolic amino acid concentrations were in the range of those reported in human muscle ^33^ and similar to concentrations observed here in A549 and U87-MG cells. The simulation correlated well with the experimental data at all time points, reaching Pearson’s R of 0.97 at t=0.25h (Fig. 6G) and t=2h (Fig. 6H). Significant increases were observed for all supplemented amino acids (Arg, Gln, His, Ile, Leu, Lys, Phe, Thr, Val) within 15 min (Fig. S6E), in myotubes and in the simulation (Pearson’s R = 0.85), while alanine, serine and asparagine were the major efflux substrate. The dynamics of the system were also replicated well for media inducing inflammation in muscle (sarcopenic muscle model) (Fig. S6F). The myotube experiments suggest that concentration transients of cellular amino acids can be simulated based on plasma amino acid changes and mRNA expression data. Importantly, the data demonstrate that the relationship of extracellular and intracellular amino acid concentrations is determined by transport processes.

## Discussion

Transporters are typically classified as uniporters, symporters and antiporters. Our results suggest a more functional classification based on their role in a cellular context. To understand amino acid homeostasis in mammalian cells four different types of transporters are required, namely harmonizers, loaders, controllers and rescue transporters (Fig. 7).

**Fig. 7.**
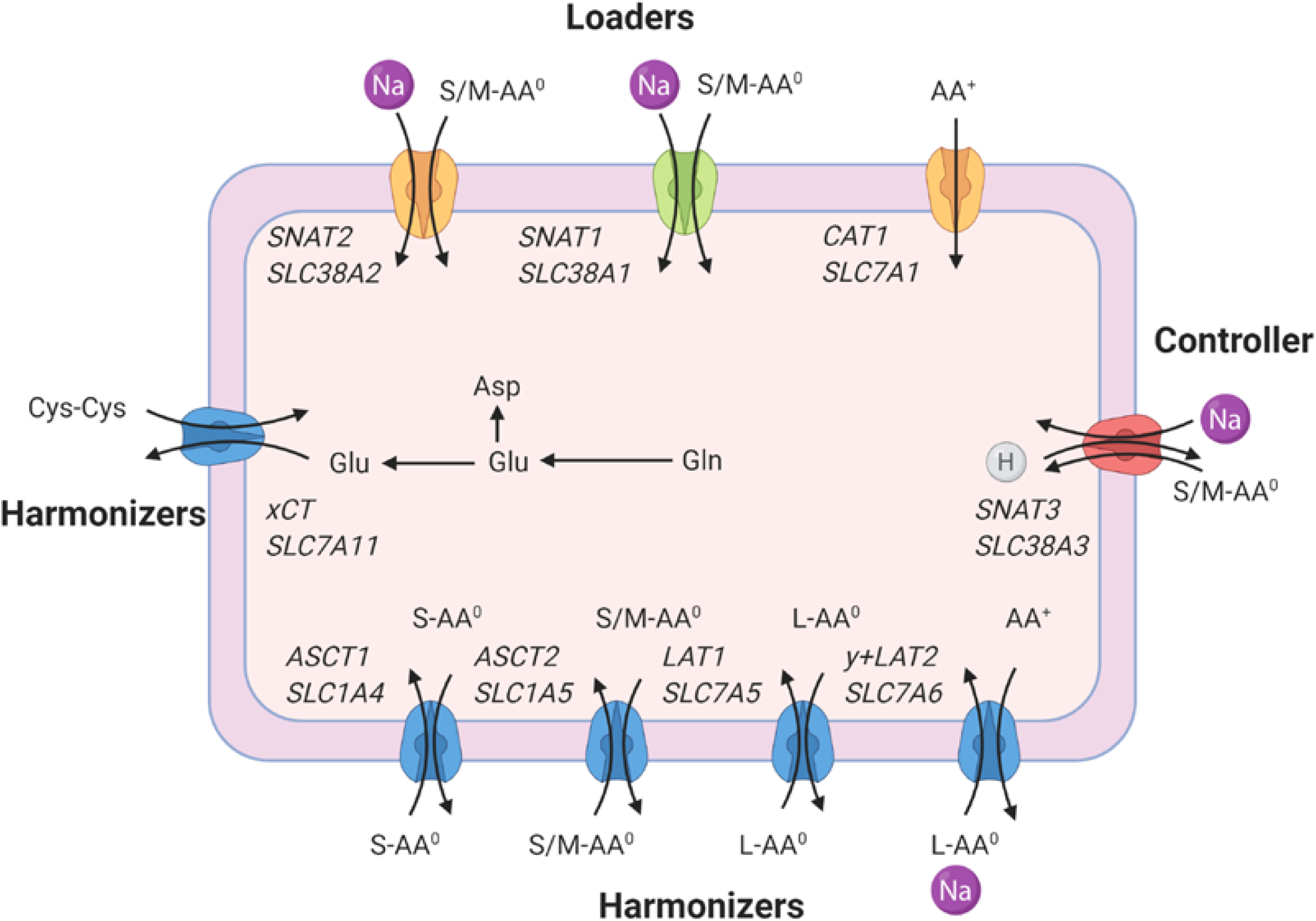
Common amino acid transporters in mammalian cells. Transporters are coloured by function. Loaders that mediate net uptake into cells are shown in green and orange. Some of these can be upregulated during amino acid starvation for rescue purposes (orange). Harmonizers that exchange amino acids are shown in blue. Controller transporters that protect against excessive accumulation are shown in red. The common transporter names and SLC acronyms are shown in italics. Amino acid (AA) charge is indicated as AA^0^, AA^-^ or AA^+^. Amino acid size is indicated as small (S), medium (M) or large (L). The most common transport reaction is indicated by the arrows, but all processes are reversible. Dominant metabolic conversions are shown, the three-letter code is used for individual amino acids. Cys-Cys is cystine.

(i) Harmonizers are the main contributors to a cell’s amino acid transport activity. These are rapid antiporters for a group of amino acids such as large amino acids (LAT1) or polar amino acids (ASCT2). In order to maintain a harmonized mix of all 20 proteinogenic amino acids, these transporters are significantly faster than net uptake. Consistently, LAT1 is the dominant transporter for large neutral amino acids, while ASCT2 dominates the transport of small and polar amino acids. The harmonizer y^+^LAT2 forms the nexus between neutral and cationic amino acids. Although it was initially thought that its main function was to efflux cationic amino acids, as observed in its close homologue y^+^LAT1 in the kidney and intestine ^34^, modelling indicates that it can mediate uptake of cationic amino acids. This has also been observed in some cell types ^35^, but the resulting cytosolic concentrations are low. As a result, not all cell lines require expression of the canonical cationic amino acid transporter CAT-1 ^10^. Antiporters show significant asymmetry with cytosolic K_M_-values being 100-1000-fold higher than extracellular K_M_-values ^36-38^.

(ii) Amino acid loaders accumulate amino acids in the cytosol. SNAT1, for instance, mediates the net uptake of a group of amino acids such as small and polar neutral amino acids ^11^. Loaders must have at least one overlapping amino acid substrate with the harmonizers, but typically share several substrates. Glutamine and alanine are ideal substrates as they are highly abundant in blood plasma and are major substrates for SNAT1. SNAT1 is a Na^+^/neutral amino acid symporter using the electrical and chemical driving force of Na^+^ to accumulate its substrates in the cytosol as shown in equation 5^11^:

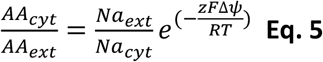

Under physiological conditions, this allows for an approx. 100-fold accumulation inside the cytosol. In conjunction with harmonizers, SNAT1 is able to lift the concentration of all neutral amino acids above those observed in the blood plasma or cell culture media. Over time, SNAT1 would accumulate all neutral amino acids to the same ratio as its own substrates.

(iii) Transporters that lower the accumulation of amino acids (controllers) to avoid excessive amino acid accumulation by loaders. Several transporters fall into this category, such as SNAT3/5, LAT3/4. For instance, SNAT3/5 mediate Na^+^-amino acid symport/H^+^-antiport. The transport process is therefore electroneutral, and accumulation of amino acids only depends on the prevailing Na^+^ and H^+^ gradients ^39^ allowing for a modest 15-fold accumulation as shown in equation 6.

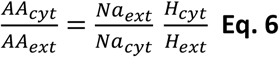

This is significantly less than the 100-fold accumulation generated by SNAT1. Accordingly, modelling shows that SNAT5 is a significant efflux pathway for highly abundant amino acids, such as alanine, glycine and glutamine. Notably, the substrate affinity of these transporters is very low. As a result, efflux only becomes considerable when intracellular substrates concentrations reach several millimolar. Additional mechanisms to avoid excessive accumulation of amino acids are metabolism and trans-inhibition of amino acid loaders, such as SNAT1/2 ^40^.

The final group (iv) are rescue transporters that can be actively up-regulated when amino acids are depleted. SNAT2 for instance is highly regulated at the transcriptional, translational and protein level ^41^. Accordingly, we could only detect marginal activity of SNAT2 in well-nourished cells ^18^. CAT1 is also actively regulated at multiple levels ^42^, allowing for the active import of cationic amino acids. Due to the opposing fluxes between CAT1 and y^+^LAT2, cationic amino acid levels in A549 and U87-MG cells were only 2-fold higher than plasma amino acid concentrations.

The general view that most amino acid transporters have indeed been identified, was supported in this study by a complete deconvolution of amino acid transport in two cell lines of very different origin, namely lung and brain. One difficulty in the deconvolution process was the presence of high-capacity substrates (leucine, alanine and glutamine) and low-capacity substrates (glutamate, arginine, glycine, proline). Standard deviations of individual incubations of the high-capacity substrates were as large as the total transport of some of the low-capacity substrates. As a result, minor components to the flux of high capacity substrates may be overlooked, because of a lack of statistical significance. However, the set of experiments we used provides significant redundancy. In general, we found that the use of canonical high-affinity substrates for each transporter was ideal to estimate V_max_.

A consistent observation was the very low activity of glutamate transporters. In the brain, extracellular glutamate levels are below 1 μM, while cytosolic levels are 5-10 mM ^43^. In this environment, glutamate transporters can work close to equilibrium, while glutamate concentrations in blood plasma and cell culture media are ∼100 µM. This poses an osmotic threat to cells in a high glutamate environment. Consequently, cancer cells express EAATs at very low levels and even release glutamate in exchange for cystine via xCT. While the intracellular affinity of xCT is unknown, it is likely to be very asymmetric to allow accumulation of close to 10 mM glutamate in the cytosol. This is corroborated by experimental observations of highly asymmetric K_M_-values in several antiporters ^36,38,44^. Cancer cells generate glutamate and aspartate via glutaminolysis ^24,45,46^ providing precursors for nucleobase synthesis or to synthesise asparagine, which can be used as an exchange substrate for antiporters, such as ASCT2 ^47^.

The results shown here have implications for amino acid signalling. Sestrin2 and CASTOR1/2 are considered sensors for leucine and arginine, respectively ^17^. Sestrin binds to leucine with a K_d_ of 20 μM ^48^ and castor binds arginine with a K_d_ of 30 μM ^49^. As shown here, cytosolic equilibrium concentrations of leucine are several-fold higher than in blood plasma, typically reaching 1-3 mM. The mechanism by which sestrin or CASTOR modulate mTORC1 may therefore include non-agonistic competition with other amino acids or other mechanisms which modify the apparent affinity of these sensors. The lysosomal arginine sensor SNAT9 ^50-52^ has lower amino acid affinity, but amino acid levels in lysosomes are unknown ^17^. It is worth noting that several amino acid transporters such as SNAT2, SNAT9 and PAT4 are considered transceptors and shown to regulate mTORC1 independent of their transport function ^53,54^. Structural evidence for the transceptor concept has recently emerged ^55^. Amino acid signalling interacts with the amino acid transportome. GCN2 signalling increases expression of loaders and harmonizers, such as SNAT2 ^18^, xCT, ASCT1, ASCT2 ^56^ and CAT1 ^57^. mTORC1 also appears to increase the transcription of a subset of ATF4 activated genes, including amino acid transporters xCT, LAT1, CAT1, ASCT2, ASCT1 and SNAT1 ^58^.

We have shown here that amino acid transporters play a key role in maintaining elevated pools of amino acids within the cytosol and establish a relationship between plasma and intracellular amino acid levels, which has been elusive heretofore ^17^. The methodology can be extended to include organellar transport processes, such as lysosomal and mitochondrial amino acid transport if volume data are available and transporters are similarly well characterised as plasma membrane transporters. Moreover, it can be used to model other types of transport in mammalian cells, such as drug transport. For the first time, this allows for a systems-level understanding of amino acid homeostasis in mammalian cells.

## Acknowledgements

The development of JDFC went through many earlier versions using different platforms. The authors would like to thank Jue Sheng Ong (QIMR Berghofer Medical Research Institute) and Ben Corry (ANU Research School of Biology) for the initial attempts to develop transport simulation and Avinash Upadhya and Woei Ming (Steve) Lee (ANU Research School of Engineering) for considerable help with Matlab scripting. Adam Carroll and Thy Truong (ANU College of Science Joint Mass Spectroscopy facility) helped with developing the amino acid analysis. The authors thank Axcella Health team members Murat Cokol and Shuran Xing for their help with data modelling and statistical analysis. We thank Michael Hamill and Meredith Duffy (Axcella Health) for their enthusiastic support of this study. We are indebted to Lon J. Van Winkle (Rocky Vista University) and Reinhard Krämer (University of Cologne) for commenting on drafts of the manuscript. Work associated with this project was funded by Australian Research Council Grant DP180101702.

## Author contributions

Conceptualization: S.B.; Methodology: G. G-C, K.J.; Software: J.V., Z.Z.; Validation: G. G-C, K.J., A.B., W.C.C.; Formal analysis: W.C.C.; S.B., G. G-C., J.V.; Investigation: G. G-C., A.B., W.C.C., K.J.; Writing: G. G-C. and S.B.; Editing: all authors; Supervision: S.B.; Project administration: S.B., B.C., Funding acquisition: S.B.

## Availability of data

The code for JDFC is available through Github on request.

## Methods

File: Materials and Methods

## Supplemental information

File: Supplemental Tables

File: Supplemental Figures

## Materials and Methods

**Table M1:**
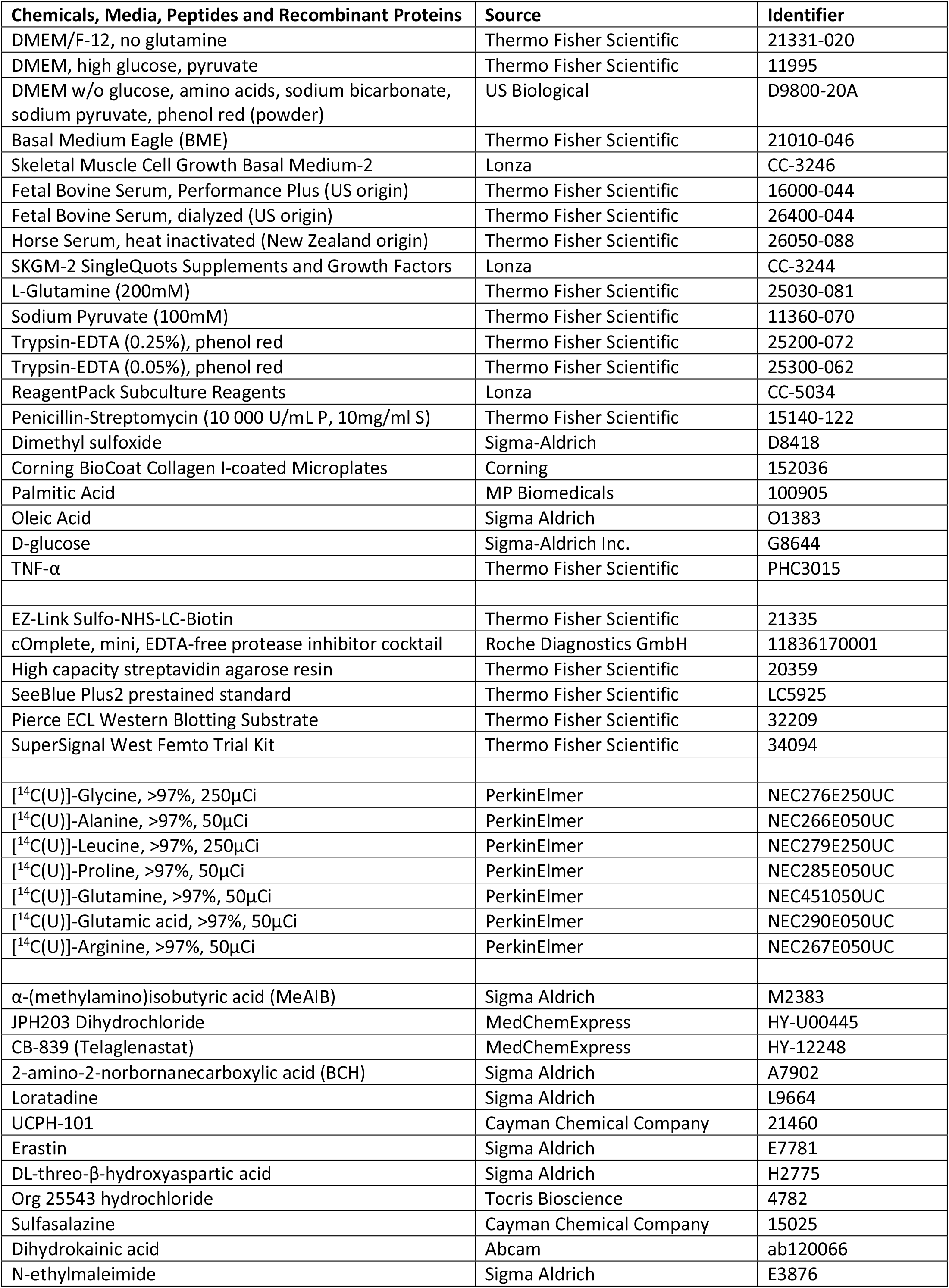
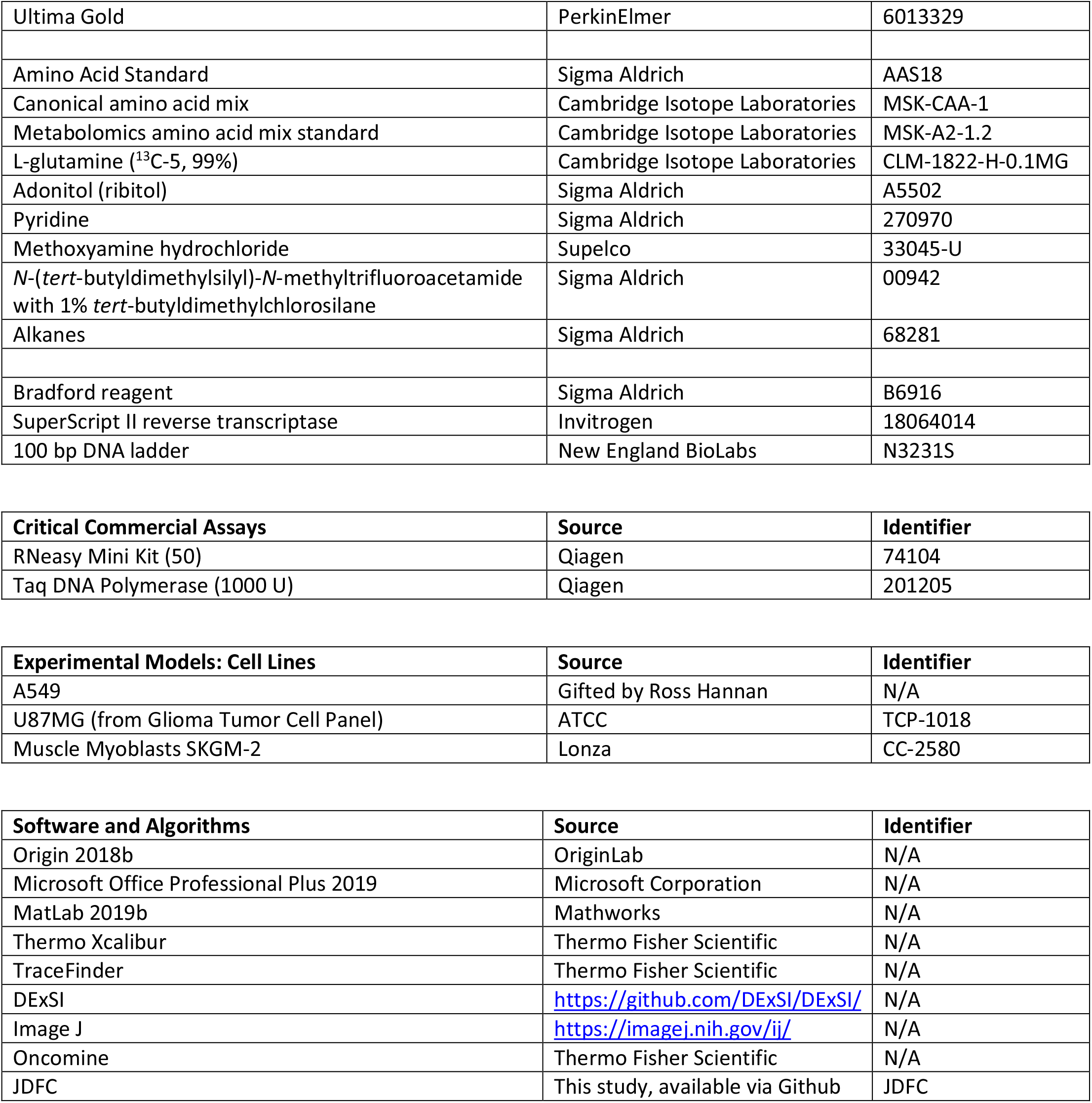
Chemicals, Media, Peptides and recombinant proteins

### Cell culture

Human lung carcinoma A549 cells were kindly gifted by Ross Hannan (John Curtin School of Medical Research, Australian National University) and human glioblastoma U87MG cells were purchased from ATCC. A549 cells were cultured in DMEM/F-12 supplemented with 10% Fetal Bovine Serum (FBS), 2mM glutamine, 10U/mL penicillin and 10µg/mL streptomycin. U87MG cells were cultured in BME containing 10% FBS, 0.5mM glutamine, 1mM sodium pyruvate, 10U/mL penicillin and 10µg/mL streptomycin (all obtained from Gibco) and non-essential amino acids at concentrations described in Bröer et al., 2016.

Primary human myoblasts (PHMs) from one healthy human donor (see Table M2) were previously screened and validated based on-post thaw viability and myotube growth. On day 0, qualified PHMs were thawed and plated in a culture flask with Skeletal Muscle Cell Growth Basal Medium-2 (SKBM-2; Lonza) fortified with SingleQuots™ supplements and growth factors (Lonza). On day 1, PHMs were collected with subculture reagents (Lonza) and seeded onto Collagen I-coated 96-well plates (Corning) at a density of 3200 cells/well.

On day 2, the medium was replaced with DMEM supplemented with 2% horse serum and 1% penicillin-streptomycin (all obtained from Gibco) to achieve myotube differentiation, and the cells were incubated at 37°C for five days prior to experimentation.

**Table M2.** Primary cells donor sourcing and information

| Donor ID | Cell Type | Vendor/Source | Sex | Race | Age | BMI |
| --- | --- | --- | --- | --- | --- | --- |
| HSMM | Muscle Myoblasts SkGM-2 | Lonza | Male | Caucasian | 29 | 35.7 |

To study the effects of LIVRQNacHKFT (AXA2678 constituents) on the intracellular amino acid levels in an *in vitro* myosteatosis model, differentiated PHMs grown on 96-well plates were washed thrice with amino acid-free DMEM and incubated in DMEM containing amino acid concentrations that matched those found in healthy human plasma (see Table M2). For lipotoxic stimulation, this same medium was used with the addition of 200µM free fatty acids (2:1 oleate:palmitate), 19µM L-glucose, and 10ng/mL TNF-α (Thermo Fisher). After 24 hours of equilibration in these media, cellular metabolites were extracted to measure baseline amino acid levels. In parallel, cells were incubated with the constituent amino acids of LIVQNacHKFT at specified fold concentrations above plasma levels (see Table M3). *N*-acetylcysteine (Nac) is not endogenous in plasma and was proportionally scaled to be 0.2mM or 0.5mM. Following this treatment, cellular metabolites were extracted at 0.25-, 0.5-, 1- and 2-hour time points for amino acid analysis.

**Table M3.**
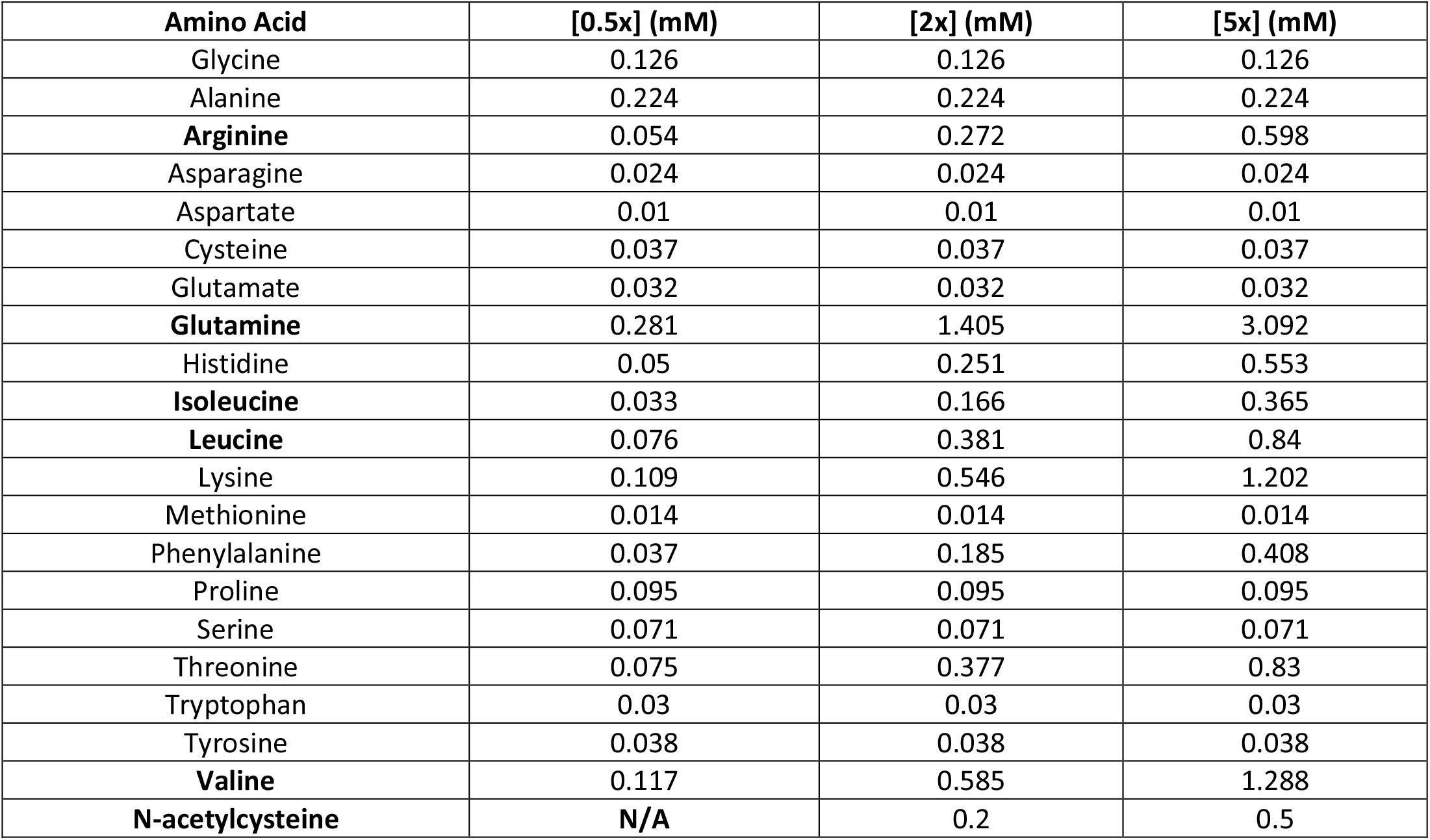
Amino acid concentrations used for PHM treatment. PHM media baseline AA concentrations (0.5x).Concentrations match mean physiological levels found in human plasma (values published in the Human Metabolome Database, HMDB; (Wishart et al., 2007). LIVRQNacHKFT (bold) were added at specified fold concentrations above plasma level (2x/5x; Nac is not endogenous in plasma and was proportionally scaled between 0.2/0.5 mM)

| Amino Acid | [0.5x] (mM) | [2x] (mM) | [5x] (mM) |
| --- | --- | --- | --- |
| Glycine | 0.126 | 0.126 | 0.126 |
| Alanine | 0.224 | 0.224 | 0.224 |
| **Arginine** | 0.054 | 0.272 | 0.598 |
| Asparagine | 0.024 | 0.024 | 0.024 |
| Aspartate | 0.01 | 0.01 | 0.01 |
| Cysteine | 0.037 | 0.037 | 0.037 |
| Glutamate | 0.032 | 0.032 | 0.032 |
| **Glutamine** | 0.281 | 1.405 | 3.092 |
| Histidine | 0.05 | 0.251 | 0.553 |
| **Isoleucine** | 0.033 | 0.166 | 0.365 |
| **Leucine** | 0.076 | 0.381 | 0.84 |
| Lysine | 0.109 | 0.546 | 1.202 |
| Methionine | 0.014 | 0.014 | 0.014 |
| Phenylalanine | 0.037 | 0.185 | 0.408 |
| Proline | 0.095 | 0.095 | 0.095 |
| Serine | 0.071 | 0.071 | 0.071 |
| Threonine | 0.075 | 0.377 | 0.83 |
| Tryptophan | 0.03 | 0.03 | 0.03 |
| Tyrosine | 0.038 | 0.038 | 0.038 |
| **Valine** | 0.117 | 0.585 | 1.288 |
| **N-acetylcysteine** | **N/A** | 0.2 | 0.5 |

### RT-PCR

Total RNA was isolated from cell lines using the RNeasy Mini Kit (Qiagen). First strand cDNA was synthesized using 2µg RNA and SuperScript II reverse transcriptase (Invitrogen) according to the manufacturer’s recommendations. PCR was performed over 30 cycles, using the *Taq* DNA polymerase kit (Qiagen) with primers listed in Table M4.

**Table M4:**
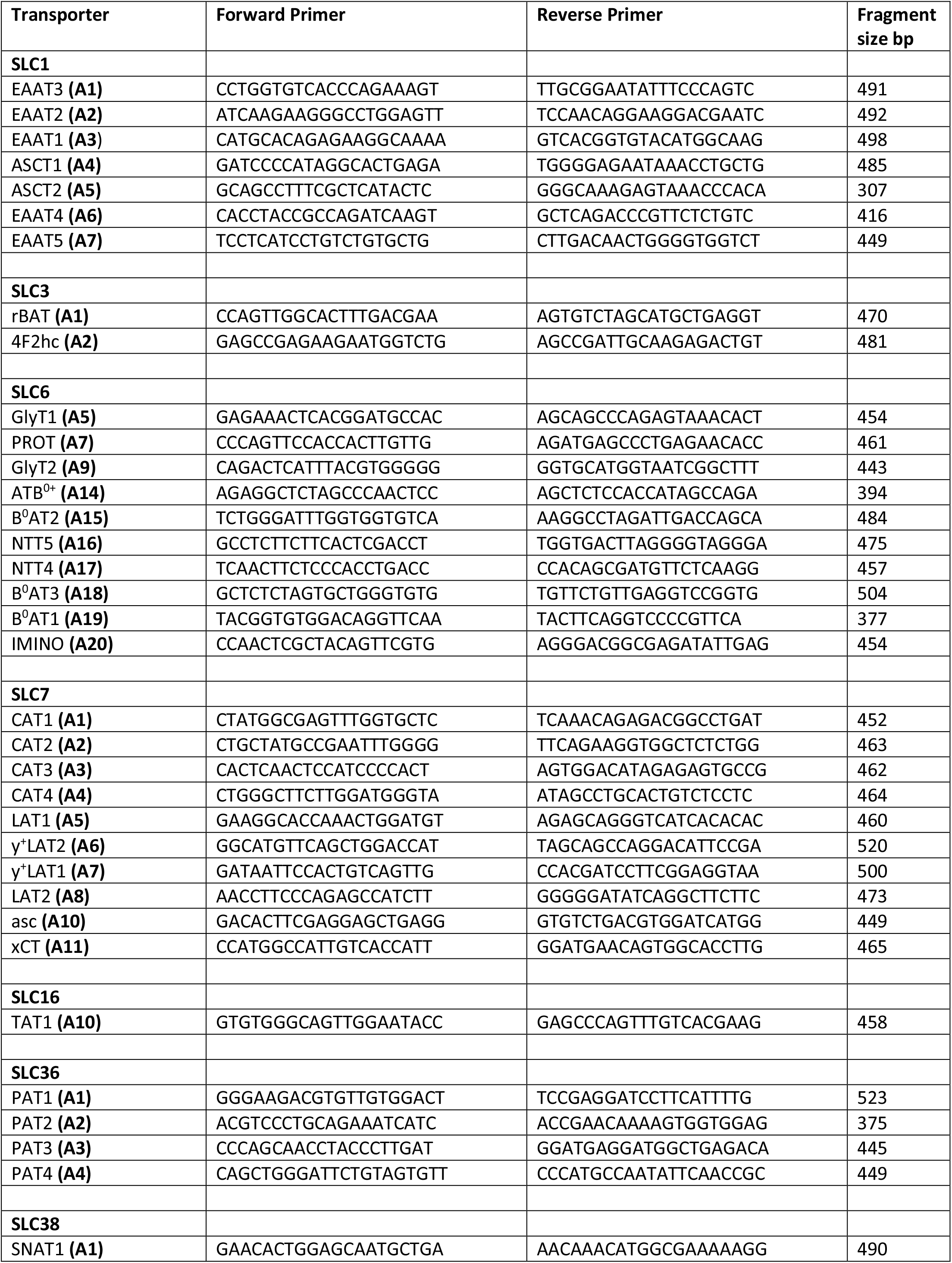
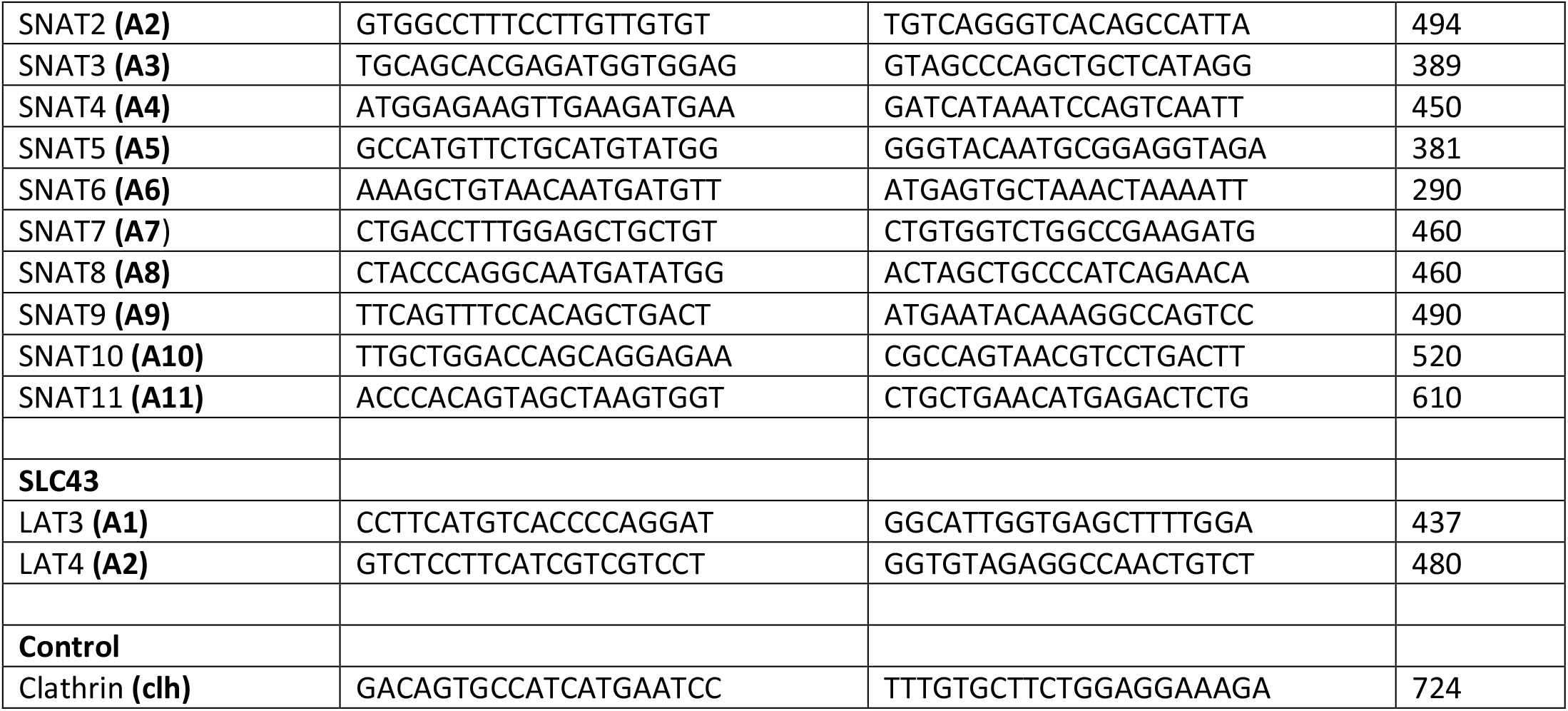
RT-PCR primers used in this study.

### RNA silencing

Silencing of B^0^AT2 in U87-MG cells was performed using Ambion Silencer Select pre-designed siRNA ID: s224342, according to the methods outlined in Bröer et al., 2016. Universal Negative Control (Sigma Aldrich, SIC001) was used as a scramble control.

### Surface biotinylation and western blotting

For surface biotinylation, cells were washed three times with modified PBS (0.6mM MgCl_2_, 1mM CaCl_2_. pH 8.0) and subsequently incubated with 1mg/mL Sulfo-NHS-LC-Biotin (Thermo Fisher) in modified PBS for 30 minutes. Excess biotinylating agent was quenched and removed with three washes of PBS supplemented with 0.1M glycine and cells were homogenized using a lysis buffer (150mM NaCl, 20mM Tris, 1% Triton X-100, pH 7.5) containing protease inhibitors (Roche). Lysates were incubated for 2h on ice with occasional mixing and then cleared by centrifugation. Protein concentration was determined by the Bradford assay to normalize the mass of protein added to high capacity streptavidin agarose resin (Thermo Fisher). After overnight incubation with slow rotation at 4°C, unbound protein was removed by five washing cycles of the resin with lysis buffer. Biotinylated proteins were mixed with 4xLDS sample buffer and were disassociated from the resin using sample reducing reagent (Thermo Fisher) and by incubation at 90°C for five minutes. Proteins were then separated on 4-12% Bolt Bis-Tris gels (Thermo Fisher) for 50 minutes and transferred to a nitrocellulose membrane (GE Healthcare). Membranes were blocked in 5% (w/v) skim milk-powder in PBS with 0.15% Tween 20 overnight, before being incubated with primary and secondary antibodies as listed in Table M5. Densitometry of protein bands was performed using Image J (Schneider et al., 2012).

**Table M5:**
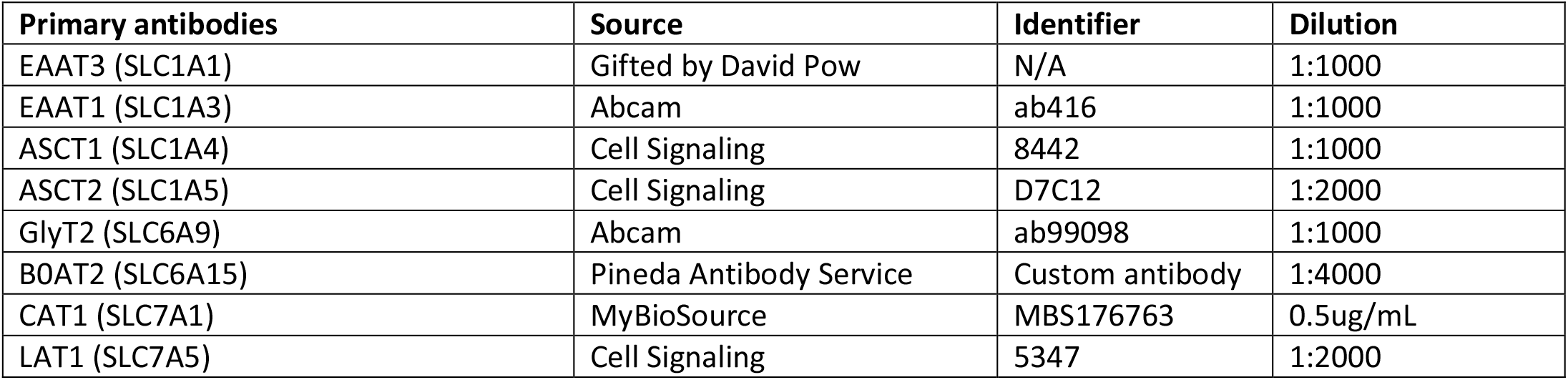
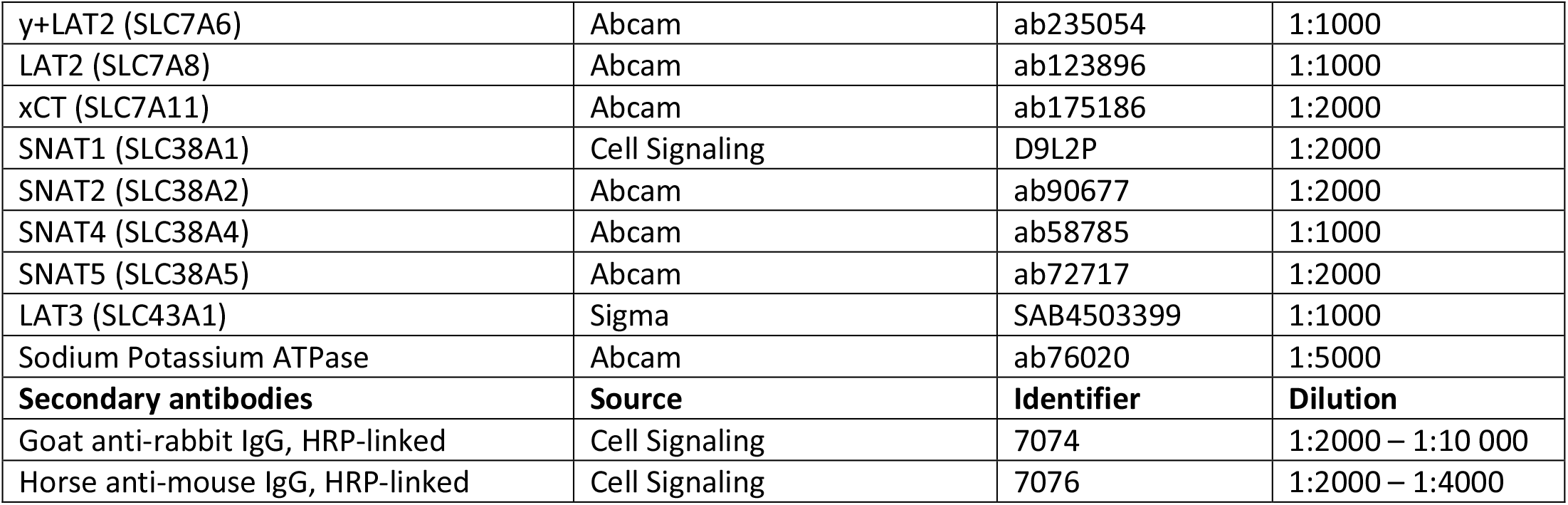
Primary and secondary antibodies used in this study

### Amino acid flux assays

Amino acid transport velocities of cells were determined by measuring the uptake of radiolabeled amino acids during the linear range of accumulation (two minutes for leucine, alanine and glutamine, and six minutes for all other substrates). Experiments were conducted on cells grown to confluence in 35 mm dishes at 37°C. A combination of Hanks’ Balanced Salt Solution (HBSS; 137mM NaCl; 5.4mM KCl; 2.7mM Na_2_HPO_4_; 1mM CaCl_2_, 0.5mM MgCl; 0.44mM KH_2_PO_4_; 0.4mM MgSO_4_; 5mM D-glucose, 5mM HEPES; pH7.4) and HBSS where sodium was substituted with lithium or *N*-methyl-*D*-glucamine (NMDG), were used to discriminate between transporters. Amino acid transporter inhibitors and competing substrates were also used to further parse the activities of transporters with similar functional characteristics.

Briefly, cells were washed with the appropriate HBSS solution prior to adding HBSS containing a given amino acid in radiolabeled (all purchased from PerkinElmer, ≥2000 cpm/nmol) and unlabeled (100µM or 300µM) forms, and in the presence or absence of inhibitors and competing substrates. After two or six minutes, cells were washed with ice-cold HBSS and lysed with 0.1M HCl. Cells were scraped and homogenized, with one portion of the homogenate transferred to a scintillation vial for counting while the other was used for protein determination to normalize the rates of uptake. Concentrations of inhibitors and incubation time are listed in Table S1 (Supplementary material).

### Cell volumetry

The volumes of A549 and U87MG cells were measured using a Multisizer 4 (Beckman Coulter) with a 100µm aperture tube. Prior to volume measurement, cells were trypsinized, pelleted and resuspended in calcium-free HBSS. This same solution was used as the electrolyte solution for the aperture tube. Approximately 5 × 10^5^ cells were diluted into ten milliliters of calcium-free HBSS in a cuvette. Cell volume measurements were taken over 60 seconds, and distributed in 200 bins spanning a volume range of 500fL to 6000fL. A lognormal fit was applied to the volume distribution data to determine an approximation of the mean cell volume for each cell line.

### Amino acid equilibration experiments and LC-MS

To study changes in cellular amino acid equilibria, A549 and U87MG cells were incubated in supplemented BME or a modified HBSS solution (HBSS supplemented with 23mM NaHCO_3_) with all twenty proteinogenic amino acids at a baseline concentration of 100µM. Cells were initially seeded at approximately 200 000 cells per 35 mm dish. Once confluent (two days post-seeding for A549 cells and five days for U87MG cells), cells were washed and incubated in the media for 30 minutes at 37°C. Each dish was then washed once with ice-cold 0.9% NaCl solution followed by a second wash with ice-cold milli-Q water, before immediately snap freezing the cells in liquid nitrogen, adding 600µL of extraction solution (60:40 methanol:water) with isotopically-labelled amino acid internal standards (MSK-A2-1.2 and CLM-1822-H-0.1, Cambridge Isotope Laboratories, all at 1.5µM), and placed on dry ice. Cells were then thawed, collected and mixed with an equal volume of chloroform. The lysate was vortexed for five minutes followed by another five minutes of sonication and the aqueous phase was cleared by centrifugation, transferred to a new microcentrifuge tube and placed in a vacuum concentrator for three hours. Metabolites were resuspended in acetonitrile:10mM ammonium acetate (90:10 + 0.15% formic acid) for LCMS analysis. This method was also used to measure glutaminase activity in A549 and U87MG cells using CB-839, with the exception of the incubation time being extended to 60 minutes.

Amino acids were analyzed with an Orbitrap Q-Exactive Plus coupled to an UltiMate 3000 RSLCnano system (Thermo Fisher). Separation of analytes was achieved using a 3µm ZIC cHILIC 2.1 × 150mm column (EMD Millipore) with a gradient elution of two mobile phases over 21 minutes at a flow rate of 0.4mL/min. Solvent A was comprised of 10mM ammonium acetate in water with 0.15% formic acid and solvent B consisted of acetonitrile with 0.15% formic acid. Solvent A was held at 10% from 0 to 6.0 min, linearly increased to 15% between 6.0 and 6.1 min, increased again to 26% from 6.1 to 10.0 min and then to 36% from 10.0 to 12.0 min and to 64% from 12.0 to 12.1 min where it was held until 16.0 min. From 17.0 to 17.1 min, solvent A was decreased to 10% and held until 21 min. Analytes were ionized in positive mode, and analyzed with a combination of survey scans (73-194 *m/z*, R=35 000) and parallel reaction monitoring (R=17 500), both with a target of 5 × 10^4^ ions, a maximum injection time of 50ms and an isolation window of 0.5 *m/z*. Collision energies were optimised for each analyte (see Table M5). Quantification was achieved using Xcalibur (Thermo Fisher) by integrating the base peak in the MSMS spectra of all amino acids and their isotopes, except glycine, alanine and aspartate (due to the production of daughter ions that were below the *m/z* instrument detection threshold or unreliable fragmentation patterns) for which parent ion peaks were selected for quantification (see Table M5). Tryptophan and asparagine could not be quantified due to the absence of their respective isotopes in the internal standards mixture and cysteine was omitted from analysis due to its high intrinsic reactivity. Cytosolic concentrations were determined according to the following equation:

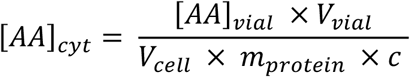

Where *V_cell_*is the mean cell volume, *m_protein_*is the mean protein mass per dish measured in satellite dishes in each experiment, and *c* is the correlate of protein mass and cell number (7036 ± 488 cells/µg for A549 cells and 7092 ± 830 cells/µg for U87-MG cells). This correlate was determined by trypsinizing, pelleting and resuspending A549 and U87-MG cells in HBSS, measuring cell density with a hemocytometer and measuring protein mass using the Bradford assay.

**Table M5:** Parameters of amino acid quantification

| Analyte | Molecular formula | MW (g mol<sup>-1</sup>) | Parent ion (m/z) | CE (eV) | T<sub>R</sub> (min) | Quantification | Base peak product ion (m/z) | Qualifier product ion (m/z) | LOD (ppm) | LLOQ (ppm) | Linear regression equation | Correlation coefficient | Ionisation polarity |
| --- | --- | --- | --- | --- | --- | --- | --- | --- | --- | --- | --- | --- | --- |
| Glycine | C<sub>2</sub>H<sub>5</sub>NO<sub>2</sub> | 75.07 | 76.0393 | - | 12.57 | a | - | - | 13.628 | 22.713 | y=0.08387x +0.01813 | 0.99958 | + |
| Alanine | C<sub>3</sub>H<sub>7</sub>NO<sub>2</sub> | 89.09 | 90.0550 | - | 11.95 | a | - | - | 15.02 | 25.034 | y=0.07653x – 0.03334 | 0.99920 | + |
| Valine | C<sub>5</sub>H<sub>11</sub>NO<sub>2</sub> | 117.15 | 118.0863 | 10.0 | 10.05 | b | 72.0813 | 72.0813 | 0.129 | 0.215 | y=0.10335x +0.11064 | 0.99886 | + |
| Leucine | C<sub>6</sub>H<sub>13</sub>NO<sub>2</sub> | 131.17 | 132.1019 | 10.0 | 8.98 | b | 86.0965 | 69.0704 | 0.161 | 0.2683 | y=0.0953x +0.04235 | 0.99951 | + |
| Isoleucine | C<sub>6</sub>H<sub>13</sub>NO<sub>2</sub> | 131.17 | 132.1019 | 10.0 | 8.57 | b | 86.0965 | 86.0965 | 0.2725 | 0.4541 | y=0.09675x +0.05373 | 0.99916 | + |
| Methionine | C<sub>5</sub>H<sub>11</sub>NO<sub>2</sub>S | 149.21 | 150.0583 | 13.5 | 9.53 | b | 56.0500 | 104.0530 | 0.1516 | 0.2526 | y=0.09664x +0.02959 | 0.99975 | + |
| Phenyl-alanine | C<sub>9</sub>H<sub>11</sub>NO<sub>2</sub> | 165.19 | 166.0865 | 12.0 | 8.48 | b | 120.0811 | 120.0811 | 0.037 | 0.0617 | y=0.05395x+0.1147 | 0.99827 | + |
| Proline | C<sub>5</sub>H<sub>9</sub>NO<sub>2</sub> | 115.13 | 116.0706 | 16.0 | 10.15 | b | 70.0654 | 70.0654 | 0.0387 | 0.0645 | y=0.08993x+0.06677 | 0.99892 | + |
| Serine | C<sub>3</sub>H<sub>7</sub>NO<sub>3</sub> | 105.09 | 106.0499 | 11.0 | 12.94 | b | 60.0448 | 60.0448 | 3.5664 | 5.944 | y=0.09713x+0.22671 | 0.99778 | + |
| Threonine | C<sub>4</sub>H<sub>9</sub>NO<sub>3</sub> | 119.12 | 120.0655 | 12.0 | 12.10 | b | 74.0603 | 74.0603 | 4.1761 | 6.9601 | y=0.09751x+0.06334 | 0.99905 | + |
| Tyrosine | C<sub>9</sub>H<sub>11</sub>NO<sub>3</sub> | 181.19 | 182.0812 | 12.5 | 11.00 | b | 136.0751 | 136.0751 | 1.9891 | 3.3151 | y=0.10508x+0.08068 | 0.99806 | + |
| Glutamine | C<sub>5</sub>H<sub>10</sub>N<sub>2</sub>O<sub>3</sub> | 146.14 | 147.0764 | 13.0 | 12.48 | b | 84.0450 | 102.0548 | 4.4133 | 7.3554 | y=0.09132x+0.11506 | 0.99604 | + |
| Aspartic acid | C<sub>4</sub>H<sub>7</sub>NO<sub>4</sub> | 133.11 | 134.0448 | - | 13.30 | a | - | - | 32.093 | 53.488 | y=0.09707x+0.00648 | 0.99952 | + |
| Glutamic acid | C<sub>5</sub>H<sub>9</sub>NO<sub>4</sub> | 147.13 | 148.0604 | 12.0 | 12.75 | b | 84.0450 | 102.0547 | 14.753 | 24.588 | y=0.09298x+0.05271 | 0.99908 | + |
| Arginine | C<sub>6</sub>H<sub>14</sub>N<sub>4</sub>O<sub>2</sub> | 174.20 | 175.1190 | 18.0 | 14.05 | b | 70.0658 | 158.0926 | 8.4227 | 14.038 | y=0.10889x+0.02004 | 0.99974 | + |
| Lysine | C<sub>6</sub>H<sub>14</sub>N<sub>2</sub>O<sub>2</sub> | 146.19 | 147.1128 | 11.0 | 13.99 | b | 84.0808 | 101.1080 | 2.4048 | 4.008 | y=0.09899x+0.03442 | 0.99935 | + |
| Histidine | C<sub>6</sub>H<sub>9</sub>N<sub>3</sub>O<sub>2</sub> | 155.15 | 156.0768 | 15.5 | 14.05 | b | 110.0710 | 110.0710 | 0.4415 | 0.7358 | y=0.10406x+0.06246 | 0.99984 | + |

To extract amino acids from cultured myotubes, cells were washed once with cold PBS, and then quenched with 100µL ice cold extraction solvent (80:20, methanol:water) containing isotopically-labeled internal standards (MSK-CAA-1, Cambridge Isotope Laboratories, all at 1µM). Five wells of lysates from same treatment condition were pooled into Eppendorf tubes. Lysates were incubated at -20°C for 30 minutes, followed by at least two hours at -80°C, and cleared by centrifugation. Supernatants were transferred to new sets of Eppendorf tubes, desolvated in a speed vacuum concentrator and resuspended in 20µL of buffer composed of 42.5% acetonitrile, 57.4% water, 0.1% formic acid and 10mM ammonium formate. Samples were injected into a 1.7µm, 2.1 × 150mm Acquity UPLC BEH amide column (Waters) connected to an Ultimate 3000 UHPLC (Dionex) coupled to a Q Exactive mass spectrometer equipped with a HESI probe (Thermo Fisher). The liquid chromatography method consisted of flow rate of 0.4mL/min and a gradient elution of solvent A (95:5 acetonitrile:water + 0.15% formic acid) and solvent B (water + 0.15% formic acid) over a run time of 10 minutes per sample. Solvent A was held at 100% from 0 to 1.0 min, followed by a linear gradient to 90% solvent A between 1.0 and 3.0 min, 80% solvent A from 3.0 to 4.3 min and 60% solvent A from 4.3 to 5.6 min. Solvent B was set to 100% from 5.6 to 7.5 min and was returned to 0% between 7.5 and 10.0 min. The full scan range of the Q Exactive mass spectrometer was set at 50 to 300 *m/z* with a resolution of 70 000 at 200 *m/z*. The maximum injection time was 100ms and the automated gain control was targeted at 3 × 10^6^ ions. Amino acids were quantified using TraceFinder software (Thermo Scientific) by measuring the area under the peak for the targeted endogenous analyte in comparison to the isotopically labeled internal standard. Note that this analytical approach can detect most proteinogenic amino acids but cannot reliably measure Nac or cysteine, due to high intrinsic reactivity and instability for these analytes.

### Stable isotopic tracing of glutamine and GC-MS analysis

U87-MG cells were seeded onto 35-mm dishes and incubated in growth medium until they reached confluence, washed thrice with warm HBSS and incubated with modified HBSS (23mM NaHCO_3_) containing 1mM of [^13^C-5]L-glutamine (Cambridge Isotope Laboratories). Metabolites were extracted at 0.3-, 1-, 2- and 24-hour time points by aspirating HBSS, washing cells once with 0.9% NaCl, and depositing enough liquid nitrogen to cover the surface of each dish. Cells were quenched by the addition of 600µL of ice-cold extraction solvent (9:1 methanol:chloroform) and 10µL of internal standard (ribitol; 10mM) to each dish, and were incubated on dry ice for 10 minutes. Cells were then scraped and collected into microcentrifuge tubes and incubated on ice for five minutes. Lysates were cleared by centrifugation for five minutes at 4°C, before transferring 150µL of the supernatant into autosampler vials and dried in a speed vacuum concentrator for four hours. Quality control samples were also prepared by pooling equal volumes of each sample to monitor sample stability and analytical reproducibility throughout each batch sequence run.

Derivatization was carried out by a robotic Gerstel MPS2 multipurpose sampler in which dried extracts were incubated at 37°C for 90 minutes at 750 rpm after adding 20µL of pyridine containing 20mg/mL methoxyamine hydrochloride (Supelco). After this, 25µL of *N*-(*tert*-butyldimethylsilyl)-*N*-methyltrifluoroacetamide and 1% (w/v) *tert*-butyldimethylchlorosilane (MTBSTFA + 1% TBDMCSI; Sigma-Aldrich) was added to the samples, which were incubated for a further 60 minutes. Five microliters of a mixture of n-alkanes was also added to the samples before injection into the GC-MS instrument.

One microliter of each sample was injected in split and splitless mode in an Agilent 7890A gas chromatograph coupled to an Agilent 5975C single quadrupole mass spectrometer. The Agilent 7890A was equipped with a Varian factor four VF-5 capillary column (30m long, 0.25mm inner diameter and 0.25µm film thickness). The injector temperature was set to 230°C and helium was used as a carrier gas at a flow rate of 1mL/min. Oven temperature was initially was held at 100°C for one minute and increased to 270°C at rate of 7°C per minute, then rising to 300°C at a rate of 10°C per minute and holding at this temperature for one minute. The total run time was therefore 29 minutes per sample. A solvent delay was also added to the method until 7.1 minutes. The electron impact ion source was kept at 260°C and filament bias at -70eV. The mass spectrometer was operated in full scan mode ranging from 40-600 *m/z* using a scan rate of 3.6Hz.

The fractional labelling of the carbon atoms was determined from the mass isotopomer distributions of the following fragments: malate (*m*/*z* 233, C1–C4), aspartate (*m*/*z* 302, C1–C4), glutamate (*m*/*z* 432, C1–C5) and glutamine (*m*/*z* 431, C1–C5). The mass isotopomer distributions were corrected for natural abundance using the program Data Extraction for Stable Isotope-labelled metabolites (DEXSI) software package (Dagley and McConville, 2018).

