## Supplemental tables for "A unified model of amino acid homeostasis in mammalian cells"

### Table S1: Experimental conditions to isolate amino acid transport activities.

| Transport Substrate | Ion | Inhibitor/competitor (conc.) | Detectable Transporters |
|---|---|---|---|
| Arginine. | Na+ | None | ATB$^{0,+}$ / CAT1,2,3 / y+LAT1,2 / b$^{0,+}$AT |
| 100 µM. | Na+ | Me-DL-Trp (2.5 mM, ATB$^{0,+}$) | / CAT1,2,3 / y+LAT1,2 / b$^{0,+}$AT |
| 6 minutes. | Na+ | NEM (0.4 mM pre-treated) | ATB$^{0,+}$ / / y+LAT1,2 / b$^{0,+}$AT |
| | Na+ | Leucine (1 mM) | / CAT1,2,3 / / |
| | NMDG | Leucine (1 mM) | / CAT1,2,3 / y+LAT1,2 / |
| | Na+ | Arginine (10 mM) | None |
| Glutamate | Na+ | None | EAAT1,2,3,4,5 / xCT |
| 25 µM. | NMDG | None | / xCT |
| 6 minutes. | Na+ | Cysteine (10 mM) | EAAT1,2, ,4,5 / xCT |
| | Na+ | Dihydrokainate (200 µM, EAAT2) | EAAT1, ,3,4,5 / xCT |
| | Na+ | UCPH101 (10 µM, EAAT1) | EAAT ,2,3,4,5 / xCT |
| | Na+ | TBOA (0.1 mM and 0.5 mM) | / xCT |
| | Na+ | Sulfasalazine (0.5 mM, xCT) | EAAT1,2,3,4,5 / |
| | Na+ | Glutamate (10 mM) | None |
| Leucine | Na+ | None | ATB$^{0,+}$ / B$^0$AT1,2,NTT4 / LAT1,2,3,4 / y+LAT1,2 / SNAT2 |
| 100 µM. | NMDG | None | / LAT1,2,3,4 / |
| 2 minutes. | Na+ | Me-DL-Trp (2.5mM, ATB$^{0,+}$) | / B$^0$AT1,2,NTT4 / LAT1,2,3,4 / y+LAT1,2 / SNAT2 |
| | Na+ | Cinromide (30 µM, B$^0$AT1) | ATB$^{0,+}$ / B$^0$AT 2,NTT4 / LAT1,2,3,4 / y+LAT1,2 / SNAT2 |
| | Na+ | Loratadine (50 µM, B$^0$AT2) | ATB$^{0,+}$ / B$^0$AT1 ,NTT4 / LAT1,2,3,4 / y+LAT1,2 / SNAT2 |
| | Na+ | BCH (10 mM) | / NTT4 / / y+LAT1,2 / SNAT2 |
| | Na+ | JPH203 (3 µM, LAT1) | ATB$^{0,+}$ / B$^0$AT1,2,NTT4 / LAT 2,3,4 / y+LAT1,2 / SNAT2 |
| | Na+ | Arginine (1 mM) | / B$^0$AT1,2,NTT4 / LAT1,2,3,4 / / SNAT2 |
| | Na+ | MeAIB (10 mM) | ATB$^{0,+}$ / B$^0$AT1,2,NTT4 / LAT1,2,3,4 / y+LAT1,2 / |
| | Na+ | Leucine (30 mM) | None |
| Phenylalanine | Na+ | None | ATB$^{0,+}$ / B$^0$AT1 / LAT1,2,3,4 / y+LAT1,2 / TAT1 |
| 100 µM | NMDG | None | / LAT1,2,3,4 / / TAT1 |
| 2 minutes | Na+ | NEM (0.4 mM pretreated) | ATB$^{0,+}$ / B$^0$AT1 / LAT1,2, / y+LAT1,2 / TAT1 |
| | Na+ | Me-DL-Trp (2.5 mM, ATB$^{0,+}$) | / B$^0$AT1 / LAT1,2,3,4 / y+LAT1,2 / TAT1 |
| | Na+ | Cinromide (3 µM, B$^0$AT1) | ATB$^{0,+}$ / / LAT1,2,3,4 / y+LAT1,2 / TAT1 |
| | Na+ | JPH203 (3 µM, LAT1) | ATB$^{0,+}$ / B$^0$AT1/ LAT 2,3,4 / y+LAT1,2 / TAT1 |
| | Na+ | Arginine (1 mM) | / B$^0$AT1/ LAT1,2,3,4 / / TAT1 |
| | NMDG | BCH (10 mM) | ATB$^{0,+}$ / B$^0$AT1/ / y+LAT1,2 / TAT1 |
| | +Na | Phenylalanine (30 mM) | None |
| Alanine | Na+ | None | ASCT1,2 / ATB$^{0,+}$ / B$^0$AT1,2 / asc-1 / LAT2 / PAT1,2 / SNAT1,2,4,5 |
| 100 µM | NMDG | None | / / / asc-1 / LAT2 / PAT1,2 / |
| 2 minutes | Na+ | GPNA | ASCT1, / / B$^0$AT1,2 / asc-1 / LAT2 / PAT1,2 / SNAT ,4,5 |
| | Na+ | Me-DL-Trp (2.5 mM, ATB$^{0,+}$) | ASCT1,2 / / B$^0$AT1,2 / asc-1 / LAT2 / PAT1,2 / SNAT1,2,4,5 |
| | Na+ | Cinromide (3 µM, B$^0$AT1) | ASCT1,2 / ATB$^{0,+}$ / B$^0$AT ,2 / asc-1 / LAT2 / PAT1,2 / SNAT1,2,4,5 |
| | Na+ | BMS-466442 (2uM, asc-1) | ASCT1,2 / ATB$^{0,+}$ / B$^0$AT1,2 / / LAT2 / PAT1,2 / SNAT1,2,4,5 |
| | Na+ | Loratadine (50 µM) | ASCT1,2 / ATB$^{0,+}$ / B$^0$AT1 / asc-1 / LAT2 / PAT1,2 / SNAT1,2,4,5 |
| | NMDG | BCH (10 mM) | ASCT1,2/ / / asc-1 / / PAT1,2 / SNAT1,2,4,5 |
| | Na+ | MeAIB (10 mM) | ASCT1,2 / ATB$^{0,+}$ / B$^0$AT1,2 / asc-1 / LAT2 / / SNAT 5 |
| | Na+ | GABA (10 mM) | ASCT1,2 / ATB$^{0,+}$ / B$^0$AT1,2 / asc-1 / LAT2 / PAT ,2 / SNAT1,2,4,5 |
| | Na+ | Betaine (10 mM) | ASCT1,2 / ATB$^{0,+}$ / B$^0$AT1,2 / asc-1 / LAT2 / PAT1,2 / SNAT1, ,4,5 |
| | Na+ | Alanine (30 mM) | None |
| Glutamine | Na+ | None | ASCT2 / ATB$^{0,+}$ / B$^0$AT1,2 / LAT1,2 / y+LAT1,2/ SNAT1,2,3,4,5 |
| 100 µM | NMDG | None | / / / LAT1,2 / / |
| 2 minutes. | Na+ | Me-DL-Trp (2.5 mM, ATB$^{0,+}$) | ASCT2 / / B$^0$AT1 / LAT1,2 / y+LAT1,2/ SNAT1,2,3,4,5 |
| | Na+ | Cinromide (3 µM, B$^0$AT1) | ASCT2 / ATB$^{0,+}$ / / LAT1,2 / y+LAT1,2/ SNAT1,2,3,4,5 |
| | Na+ | JPH203 (3 µM, LAT1) | ASCT2 / ATB$^{0,+}$ / B$^0$AT1 / LAT 2 / y+LAT1,2/ SNAT1,2,3,4,5 |
| | Na+ | BCH (10 mM) | ASCT2 / ATB$^{0,+}$ / / / y+LAT1,2/ SNAT1,2,3,4,5 |
| | Na+ | Arginine (10 mM) | ASCT2 / / B$^0$AT1 / LAT1,2 / / SNAT1,2,3, ,5 |
| | Li+ | None | / / / LAT1,2 / y+LAT1,2/ SNAT 3, ,5 |
| | Na+ | MeAIB (10 mM) | ASCT2/ ATB$^{0,+}$ / B$^0$AT1 / LAT1,2 / y+LAT1,2/ SNAT ,3, ,5 |
| | Na+ | Betaine (10 mM) | ASCT2 / ATB$^{0,+}$ / B$^0$AT1 /LAT1,2 / y+LAT1,2/ SNAT1, ,3,4,5 |
| | NMDG | BCH (10mM) | ASCT2 / ATB$^{0,+}$ / B$^0$AT1 / / y+LAT1,2/ SNAT1,2,3,4,5 |
| | Na+ | Glutamine (10 mM) | None |
| Glycine | Na+ | None | ASCT2/ GlyT1,2 / ATB$^{0,+}$ / B$^0$AT1 / asc-1 / LAT2 / PAT1,2 / SNAT1,2,4,5 |
| 100 µM | NMDG | None | / / / / asc-1 / LAT2 / PAT1,2 / |
| 6 minutes. | Na+ | ALX5407(1 µM, GlyT1) | ASCT2 / GlyT ,2 / ATB$^{0,+}$ / B$^0$AT1 / asc-1 / LAT2 / PAT1,2 / SNAT1,2,4,5 |
| | Na+ | ORG25543 (0.2 µM, GlyT2) | ASCT2/ GlyT1, / ATB$^{0,+}$ / B$^0$AT1 / asc-1 / LAT2 / PAT1,2 / SNAT1,2,4,5 |
| | Na+ | Me-DL-Trp (2.5 mM, ATB$^{0,+}$) | ASCT2/ GlyT1,2 / / B$^0$AT1 / asc-1 / LAT2 / PAT1,2 / SNAT1,2,4,5 |
| | Na+ | Cinromide (3 µM, B$^0$AT1) | ASCT2/ GlyT1,2 / ATB$^{0,+}$ / / asc-1 / LAT2 / PAT1,2 / SNAT1,2,4,5 |
| | Na+ | BMS-466442 (2 µM, asc-1) | ASCT2/ GlyT1,2 / ATB$^{0,+}$ / B$^0$AT1 / / LAT2 / PAT1,2 / SNAT1,2,4,5 |
| | NMDG | BCH (10 mM) | ASCT2/ GlyT1,2 / / / asc-1 / / PAT1,2 / SNAT1,2,4,5 |
| | Na+ | MeAIB (10 mM) | ASCT2/ GlyT1,2 / ATB$^{0,+}$ / B$^0$AT1 / asc-1 / LAT2 / / SNAT 5 |
| | Na+ | GABA (15 mM) | ASCT2/ GlyT1,2 / ATB$^{0,+}$ / B$^0$AT1 / asc-1 / LAT2 / PAT ,2 / SNAT1,2,4,5 |
| | Na+ | Betaine (10 mM) | ASCT2/ GlyT1,2 / ATB$^{0,+}$ / B$^0$AT1 / asc-1 / LAT2 / PAT1,2 / SNAT1, ,4,5 |
| | Na+ | Glycine (30 mM) | None |
| Proline | Na+ | None | ASCT1 / PROT / B$^0$AT1,2 / SIT / asc-1 / PAT1,2,4 / SNAT1,2,4,5 |

| | | | |
|---|---|---|---|
| 100 µM<br>6 minutes. | NMDG | None | / / / / PAT1,2,4 / |
| | Na+ | LP-403812 (10 µM, PROT) | ASCT1 / / $B^0$AT1,2 / SIT / PAT1,2,4 / SNAT1,2,4,5 |
| | Na+ | Cinromide (3 µM, $B^0$AT1) | ASCT1 / PROT / $B^0$AT ,2 / SIT / PAT1,2,4 / SNAT1,2,4,5 |
| | Na+ | Loratadine (50 µM, $B^0$AT2) | ASCT1 / PROT / $B^0$AT1, / SIT / PAT1,2,4 / SNAT1,2,4,5 |
| | Na+ | Tryptophan (1 mM) | ASCT1 / PROT / $B^0$AT1,2 / SIT / PAT1,2, / SNAT1,2,4,5 |
| | Na+ | L-trans-4-hydroxyproline (200 µM) | / PROT / $B^0$AT1,2 / SIT / PAT1,2,4 / SNAT1,2,4,5 |
| | Na+ | GABA (15 mM) | ASCT1 / PROT / $B^0$AT1,2 / SIT / PAT ,2,4 / SNAT1,2,4,5 |
| | Na+ | MeAIB (10 mM) | ASCT1 / PROT / $B^0$AT1,2 / / PAT ,4 / SNAT ,5 |
| | Na+ | Proline (30 mM) | None |

### Table S2: Amino acid transporters and their mechanisms

Transporters are listed by solute carrier family number and common name. The corresponding transport activity in cultured cells is shown in column "Activity". Substrates are shown in one-letter code or as groups with $AA^0$ indicating neutral amino acids, $AA^+$ indicating cationic amino acids, O, ornithine, Cit, citrulline, Cn, cysteine. S: symport, A: antiport, subcellular location is shown as PM, plasma membrane; V, vesicular; M, mitochondria; Ly, lysosome.

| SLC | Acronym | Substrates | Activity | Mechanism | Loc |
|---|---|---|---|---|---|
| SLC1A1 | EAAT3 | D,E | System $X_{AG}$ | S:3Na+/1H+A:1K+ | PM |
| SLC1A2 | EAAT2 | D,E | System $X_{AG}$ | S:3Na+/1H+A:1K+ | PM |
| SLC1A3 | EAAT1 | D,E | System $X_{AG}$ | S:3Na+/1H+A:1K+ | PM |
| SLC1A4 | ASCT1 | A,S,C | System ASC | Antiporter | PM |
| SLC1A5 | ASCT2 | A,S,C,T,Q,N | System ASC | Antiporter | PM |
| SLC1A6 | EAAT4 | D,E | System $X_{AG}$ | S:3Na+/1H+A:1K+ | PM |
| SLC1A7 | EAAT5 | D,E | System $X_{AG}$ | S:3Na+/1H+A:1K+ | PM |
| SLC3A1 | rBAT | Trafficking | Heavy chains of | | PM |
| SLC3A2 | 4F2hc | subunits | Heteromeric AA | | PM |
| SLC6A5 | GlyT2 | G | System Gly | S:3Na+/1Cl- | PM |
| SLC6A7 | PROT | P | Proline transporter | S:2Na+/1Cl- | V/PM |
| SLC6A9 | GlyT1 | G | System Gly | S:2Na+/1Cl- | PM |
| SLC6A14 | $ATB^{0,+}$ | All $AA^0$, $AA^+$ | System $B^{0,+}$ | S:2Na+/1Cl- | PM |
| SLC6A15 | $B^0$AT2 | P,L,V,I,M | System$B^0$ | S:1Na+ | PM |
| SLC6A17 | NTT4/$B^0$AT3 | L,M,P,C,A,Q,S,H,G | System$B^0$ | S:2Na+/1Cl- | PM/V |
| SLC6A18 | XT2/$B^0$AT3 | G,A | System Gly | S:2Na+/1Cl- | PM |
| SLC6A19 | $B^0$AT1 | All $AA^0$ | System $B^0$ | S:1Na+ | PM |
| SLC6A20 | SIT1 | P (F,L) | System IMINO | S:2Na+/1Cl- | PM |
| SLC7A1 | CAT-1 | K,R,O | System y+ | Uniporter | PM |
| SLC7A2 | CAT-2 | K,R,O | System y+ | Uniporter | PM |
| SLC7A3 | CAT-3 | K,R,O | System y+ | Uniporter | PM |
| SLC7A4 | CAT-4 | unknown | unknown | unknown | Ly |
| SLC7A5 | LAT1/4F2hc | H,M,L,I,V,F,Y,W | System L | Antiporter | PM |
| SLC7A6 | y+LAT2/4F2hc | K,R,Q,H,M,L | System y+L | S:AA0-Na+A:AA+ | PM |
| SLC7A7 | y+LAT1/4F2hc | K,R,Q,H,M,L,A,C | System y+L | S:AA0-Na+A:AA+ | PM |
| SLC7A8 | LAT2/4F2hc | All $AA^0$ (-P) | System L | Antiporter | PM |
| SLC7A9 | b0,+AT/rBAT | R,K,O,Cn | System $b^{0,+}$ | Antiporter | PM |
| SLC7A10 | Asc-1/4F2hc | G,A,S,C,T | System asc | Antiporter | PM |
| SLC7A11 | xCT/4F2hc | E,Cn | Sytem $x^-_c$ | Antiporter | PM |
| SLC7A12 | Asc-2 | G,A,S,C,T | System asc | Antiporter | PM |
| SLC7A13 | AGT1/rBAT | Cn | Cystine transporter | Antiporter | PM |
| SLC7A14 | | K,R,O | System c | Uniporter | Ly |
| SLC16A10 | TAT1 | W,Y,F | System T | Uniporter | PM |
| SLC17A6 | VGLUT2 | E | Vesicular glutamate transporter | Uniporter | V |
| SLC17A7 | VGLUT1 | E | Vesicular glutamate transporter | Uniporter | V |
| SLC17A8 | VGLUT3 | E | Vesicular glutamate transporter | Uniporter | V |
| SLC25A2 | ORC2 | K,R,H,O,Cit | Orn/Cit carrier | Antiporter | M |
| SLC25A12 | AGC1 | D,E | Asp/Glu carrier | Antiporter | M |
| SLC25A13 | AGC2 | D,E | Asp/Glu carrier | Antiporter | M |

| | | | | | |
|---|---|---|---|---|---|
| SLC25A15 | ORC1 | K,R,H,O,Cit | Orn/Cit carrier | Antiporter | M |
| SLC25A18 | GC2 | E | Glu carrier | A:OH- | M |
| SLC25A22 | GC1 | E | Glu carrier | A:OH- | M |
| SLC32A1 | VIAAT | G,GABA | Vesicular Gly/GABAtransporter | A:H+ | V |
| SLC36A1 | PAT1 | G,P,A | Proton-amino acid transporter | S:H+ | PM,Ly |
| SLC36A2 | PAT2 | G,P,A | Proton-amino acid transporter | S:H+ | PM |
| SLC36A3 | PAT3 | unknown | unknown | unknown | PM |
| SLC36A4 | PAT4 | P,W | Amino acid sensor | unknown | PM,L |
| SLC38A1 | SNAT1 | G,A,N,C,Q,H,M | System A | S:Na+ | PM |
| SLC38A2 | SNAT2 | G,P,A,S,C,Q,N,H,M | System A | S:Na+ | PM |
| SLC38A3 | SNAT3 | Q,N,H | System N | S:Na+A:H+ | PM |
| SLC38A4 | SNAT4 | G,A,S,C,Q,N,M | System A | S:Na+ | PM |
| SLC38A5 | SNAT5 | Q,N,H,A | System N | S:Na+A:H+ | PM |
| SLC38A6 | SNAT6 | Unknown | Unknown | unknown | |
| SLC38A7 | SNAT7 | Q,N,H,A,S,D | Unknown | S:Na+ | Ly |
| SLC38A8 | SNAT8 | Q,A,R,H,D | System A | S:Na+ | PM |
| SLC38A9 | SNAT9 | Q,R,N | Lysosomal transporter | S:Na+ | Ly |
| SLC38A10 | SNAT10 | Unknown | Unknown | Unknown | |
| SLC38A11 | SNAT11 | Unknown | Unknown | Unknown | |
| SLC43A1 | LAT3 | L,I,M,F,V | System L | Uniporter | PM |
| SLC43A2 | LAT4 | L,I,M,F,V | System L | Uniporter | PM |

### Table S3: Substrate Km-Values of known amino acid transporters

**All Km –values are shown in µM.**

| **SLC1 family** | |
|---|---|
| **SLC1A1 (EAAT3)** | (Arriza et al., 1994; Kanai et al., 1995; Wang et al., 2019; Watzke et al., 2000) |
| Glutamate | 62±8 |
| Aspartate | 24±2 |
| Na | 30000 (n=3) |
| K | 4000 |
| H | 0.003 (est) |
| **SLC1A2 (EAAT2)** | (Arriza et al., 1994) |
| Glutamate | 97±4 |
| Aspartate | 7±1 |
| Na | 30000 (n=3) |
| K | 4000 (est) |
| H | 0.003 (est) |
| **SLC1A3(EAAT1)** | (Arriza et al., 1994) |
| Glutamate | 48±10 |
| Aspartate | 16±1 |
| Na | 41000 (n=3) |
| K | 4000 (est) |
| H | 0.03 (est) |
| **SLC1A4 (ASCT1)** | (Arriza et al., 1993) |
| Proline | 672±185 |
| Alanine | 71±14 |
| Serine | 88±11 |
| Cysteine | 29±6 |
| Threonine | 137±19 |
| Valine | 390±8 |
| Na | 16000 |
| **SLC1A5 (ASCT2)** | (Utsunomiya-Tate et al., 1996) |
| Asymmetry of Km | $K_{Mi}/K_{Mo}$=100 (Scalise et al., 2014) |
| Alanine | 18.4 |

| | |
|---|---|
| Serine | 18.6 |
| Threonine | 20.5 |
| Cysteine | 18.8 |
| Glutamine | 23.8 |
| Methionine | 288 |
| Glycine | 361 |
| Leucine | 367 |
| Valine | 522 |
| Glutamate | 1630 |
| Na | 5700 |
| **SLC1A6 (EAAT4)** | (Fairman et al., 1995) |
| Glutamate | 2.5±0.9 |
| Aspartate | 1±0.3 |
| Na | 30000 (est) |
| K | 4000 (est) |
| H | 0.03 (est) |
| **SLC1A7 (EAAT5)** | (Arriza et al., 1997) |
| Glutamate | 64±6 |
| Aspartate | 13±5 |
| Na | 30000 (est) |
| K | 4000 (est) |
| H | 0.03 (est) |

| | |
|---|---|
| **SLC6 family** | |
| **SLC6A5 GlyT2** | (Liu et al., 1993; Lopez-Corcuera et al., 1998) |
| Glycine | 17 |
| Na | 82000 (n=3) |
| Cl | 48500 |
| **SLC6A6 (TAUT)** | (Ramamoorthy et al., 1994) |
| Taurine | 5 |
| Na | 80000 (n=2) |
| Cl | 30000 |
| **SLC6A7 (PROT)** | (Shafqat et al., 1995) |
| Proline | 6.2 |
| | 80000 (n=2) (est) |
| | 30000 (est) |
| **SLC6A9 (GlyT1)** | (Lopez-Corcuera et al., 1998) |
| Glycine | 220 |
| Na | 90000 (n=2) |
| Cl | 16000 |
| **SLC6A14 ($ATB^{0,+}$)** | |
| Amino acid $EC_{50}$ | (Sloan and Mager, 1999) |
| Isoleucine | 6 ± 1 |
| Leucine | 12 ± 2 |
| Methionine | 14 ± 1 |
| Valine | 36 ± 2 |
| Alanine | 99 ± 36 |
| Glycine | 111 ± 30 |
| Proline | >5000 |
| Serine | 43 ± 5 |
| Cysteine | 118 ± 33 |
| Asparagine | 348 ± 84 |
| Threonine | 405 ± 80 |
| Glutamine | 633 ± 62 |
| Phenylalanine | 17 ± 1 |

| | |
|---|---|
| Tryptophan | 26 ± 6 |
| Tyrosine | 92 ± 10 |
| Histidine | 76 ± 20 |
| Lysine | 100 ± 1 |
| Arginine | 104 ± 35 |
| Na | 7400 (n=2.3) |
| Cl | 920 |
| **SLC6A15 (B$^0$AT2)** | (Broer et al., 2006; Takanaga et al., 2005a) |
| Proline | 195±16 |
| Alanine | 670±92 |
| Valine | 70 (est) |
| Methionine | 40±4 |
| Leucine | 81±9 |
| Isoleucine | 58±10 |
| Phenylalanine | 1050±112 |
| Na | 16000 |
| **SLC6A17 (NTT4)** | (Zaia and Reimer, 2009) |
| Leucine | 280 |
| Methionine | 280 (est) |
| Proline | 360 |
| Cysteine | 360 (est) |
| Alanine | 360 (est) |
| Glutamine | 5200 (est) |
| Serine | 5200 (est) |
| Histidine | 5200 (est) |
| Glycine | 5200 (est) |
| Na | 23000 |
| **SLC6A19 (B0AT1)** | (Bohmer et al., 2005; Cheng et al., 2017) |
| Methionine | 1000 (est) |
| Leucine | 1100 |
| Isoleucine | 1100 (est) |
| Valine | 1100 (est) |
| Glutamine | 3200 |
| Asparagine | 3200 (est) |
| Cysteine | 4000 (est) |
| Phenylalanine | 4700 |
| Alanine | 4100 |
| Serine | 4000 (est) |
| Glycine | 11700 |
| Tyrosine | 4700 (est) |
| Threonine | 4000 (est) |
| Histidine | 3200 (est) |
| Proline | 11000 (est) |
| Tryptophan | 4700 (est) |
| Na | 21000 |
| **SLC6A20 (SIT)** | (Kowalczuk et al., 2005; Takanaga et al., 2005b) |
| Proline | 130 |
| Na | 22000 |
| Cl | 2300 |

| **SLC7 family** | |
|---|---|
| **SLC7A1** | (Wang et al., 1991) |
| Arginine | 77 |
| Lysine | 73 |
| Ornithine | 105 |

| | |
|---|---|
| **SLC7A2** | (Kavanaugh et al., 1994) |
| Arginine | 2700 (CAT2A) 38 (CAT2B) |
| Lysine | 51 |
| Ornithine | 174 |
| **SLC7A3** | (Hosokawa et al., 1997) |
| Arginine | 103 |
| Lysine | 147 |
| Ornithine | 219 |
| **SLC7A4** | Lysosomal |
| **SLC7A5 (LAT1)** | (Yanagida et al., 2001) |
| Asymmetry of Km | $K_{Mi}/K_{Mo}$ = 1000 (Meier et al., 2002) |
| Leucine | 20 |
| Isoleucine | 25 |
| Valine | 47 |
| Methionine | 20 |
| Phenylalanine | 14 |
| Tyrosine | 28 |
| Tryptophan | 21 |
| Histidine | 12 |
| Glutamine | 1640 |
| Asparagine | 2150 |
| **SLC7A6 (y+LAT2)** | (Broer et al., 2000; Chubb et al., 2006) |
| Arg | 138 |
| Lys | 138 (est) |
| His | 138 (est) |
| Leu | 236 |
| Tyr | 236 (est) |
| Trp | 236 (est) |
| Phe | 236 (est) |
| Gln | 295 |
| Met | 236 (est) |
| Cys | 500 (est) |
| Ile | 500 (est) |
| Ala | 1000 (est) |
| Na | 5000 |
| **SLC7A7 (y+LAT1)** | (Kanai et al., 2000; Pfeiffer et al., 1999) |
| Arg | 340 |
| Lys | 68 |
| His | 68 (est) |
| Leu | 44 |
| Tyr | 80 (est) |
| Phe | 80 (est) |
| Gln | 60 (est) |
| Met | 60 (est) |
| Ile | 80 (est) |
| Na | 15000 (est) |
| **SLC7A8 (LAT2)** | (Segawa et al., 1999) |
| Asymmetry of Km | $K_{Mi}/K_{Mo}$ = 200 (Meier et al., 2002) |
| Histidine | 181 |
| Tryptophan | 58 |
| Tyrosine | 36 |
| Phenylalanine | 45 |
| Valine | 124 |
| Isoleucine | 97 |
| Leucine | 119 |

| | |
|---|---|
| Methionine | 204 |
| Glutamine | 151 |
| Asparagine | 81 |
| Cysteine | 109 |
| Threonine | 69 |
| Serine | 116 |
| Alanine | 187 |
| Glycine | 265 (Low Vmax 21%) |
| **SLC7A9 (b0,+AT)** | (Mizoguchi et al., 2001) |
| Asymmetry of Km | $K_{Mi}/K_{Mo}$ = 20 (Torras-Llort et al., 2001) |
| Arginine | 108 |
| Lysine | 394 |
| Ornithine | 195 |
| Cystine | 296 |
| Alanine | 300 (est) |
| Cysteine | 100 (est) |
| Phenylalanine | 150 (est) |
| Histidine | 500 (est) |
| Isoleucine | 400 (est) |
| Leucine | 107 |
| Valine | 300 (est) |
| Methionine | 100 (est) |
| Asparagine | 500 (est) |
| Glutamine | 500 (est) |
| Serine | 150 (est) |
| Threonine | 100 (est) |
| Tryptophan | 500 (est) |
| Tyrosine | 150 |
| **SLC7A10 (asc-1)** | (Fukasawa et al., 2000) |
| Alanine | 23 |
| Glycine | 8 |
| Serine | 11 |
| Threonine | 19 |
| Cysteine | 24 |
| Valine | 112 |
| Methionine | 139 |
| Isoleucine | 160 |
| Leucine | 245 |
| Histidine | 368 |
| Phenylalanine | 464 |
| **SLC7A11 (xCT)** | (Bassi et al., 2001) |
| Glutamate | 92 |
| Cystine | 43 |
| **SLC7A12 (AGT-1)** | (Matsuo et al., 2002; Nagamori et al., 2016) |
| Aspartate | 25 |
| Glutamate | 21 |
| Cystine | 68 |

| | |
|---|---|
| **SLC16 family** | |
| **SLC16A10** | (Kim et al., 2001) |
| Tryptophan | 3720 |
| Tyrosine | 2590 |
| Phenylalanine | 7020 |

| | |
|---|---|
| **SLC36 family** | |
| **SLC36A1 (PAT1)** | (Boll et al., 2002) |
| Glycine | 7000 |
| Proline | 2800 |
| Alanine | 7500 |
| H | 0.3 |
| **SLC36A2 (PAT2)** | (Boll et al., 2002) |
| Glycine | 590 |
| Proline | 120 |
| Alanine | 260 |
| H | 0.01 |
| **SLC36A3** | Not characterised |
| **SLC36A4** | (Pillai and Meredith, 2011) |
| Proline | 3 |
| Alanine | 1480 |
| Tryptophan | 2 |
| H | 0.3 (est) |

| | |
|---|---|
| **SLC38 family** | |
| **SLC38A1 (SNAT1)** | (Albers et al., 2001; Mackenzie et al., 2003) |
| Ala | 306 |
| Ser | 306 (est) |
| Thr | 890 (est) |
| Cys | 306 (est) |
| Met | 200 (est) |
| His | 230 (est) |
| Gln | 230 |
| Asn | 306 (est) |
| Gly | 890 (est) |
| Pro | 1000 (est) |
| Na | 9500 |
| **SLC38A2 (SNAT2)** | (Yao et al., 2000) |
| Ala | 529 |
| Ser | 529 (est) |
| Thr | 1900 (est) |
| Cys | 529 (est) |
| Met | 529 (est) |
| His | 800 (est) |
| Gln | 1650 |
| Asn | 600 (est) |
| Gly | 800 (est) |
| Pro | 800 (est) |
| Na | 20000 |
| **SLC38A3 (SNAT3)** | (Broer et al., 2002; Fei et al., 2000) |
| Glutamine | 1500 |
| Histidine | 7400 |
| Asparagine | 1300 |
| Na | 15000 |
| H | 0.1 |
| **SLC38A4 (SNAT4)** | (Hatanaka et al., 2001; Sugawara et al., 2000) |
| Ala | 4200 |
| Arg | 300 ± 40 |
| Ser | 4000 (est) |
| Gly | 1600 ± 300 |
| Cys | 4000 (est) |

| | |
|---|---|
| Asn | 3000 (est) |
| Thr | 3000 (est) |
| Pro | 3000 (est) |
| Met | 3000 (est) |
| Gln | 2500 ± 500 |
| Na | 70000 |
| **SLC38A5 (SNAT5)** | (Nakanishi et al., 2001) |
| Glycine | 15200 |
| Asn | 2500 |
| Ala | 2500 (est) |
| Ser | 2500 (est) |
| Gln | 3200 |
| His | 600 |
| Na | 11000 |
| H | 0.03 |
| **SLC38A6** | Uncharacterised |
| **SLC38A7** | Lysosomal |
| **SLC38A8** | Uncharacterised |
| **SLC38A9** | Lysosomal |
| **SLC38A10** | ER/Golgi |

| **SLC43 family** | |
|---|---|
| **SLC43A1 (LAT3)** | (Babu et al., 2003) |
| Leucine | 1024 ± 31 |
| Isoleucine | 1418 ± 47 |
| Valine | 1885 ± 107 |
| Phenylalanine | 1206 ± 53 |
| Methionine | 2000 est |
| **SLC43A2 (LAT4)** | (Bodoy et al., 2005) |
| Leucine | 3733 |
| Isoleucine | 4000 (est) |
| Valine | 4000 (est) |
| Phenylalanine | 4694 |
| Methionine | 4000 (est) |

**Table S4: Equations for different types of amino acid transporters**

Preparatory calculations for all transporters:

Fractional saturation:

Exofacial (ext=extracellular):

$$F_{iext} = \left(\frac{AAi_{ext}}{Km_{appext} + AAi_{ext}}\right)$$

Endofacial (cyt=cytosolic):

$$F_{icyt} = \left(\frac{AAi_{cyt}}{Km_{appcyt} + AAi_{cyt}}\right)$$

Total saturation $F_T = \sum F_i$

$F_{AAi}=F_i/F_T$

1) **Electroneutral Uniporter: SLC16A10, SLC43A1, SLC43A2**

$J_{Tec}$=Total flux from extracellular (e) medium to cytosol (c)

$J_{Tce}$=Total flux from cytosol (c) to extracellular medium (e)

$$J_{Tec} = J_{max}(F_{Text})$$

$$J_{Tce} = J_{max}(F_{Tcyt})$$

2) **Electrogenic Uniporter (cationic AA): SLC7A1, SLC7A2, SLC7A3**

$$J_{Tec} = J_{max}(F_{Text})\left(\beta(1 - F_{Tcyt})\right)$$

$$J_{Tce} = J_{max}(F_{Tcyt})\left(\frac{1}{\beta}(1 - F_{Text})\right)$$

3) **Na$^+$-Symporter: SLC6A15, SLC6A19, SLC38A1, SLC38A2, SLC38A4**

$$J_{Tec} = J_{max}\left((F_{Text})\left(\frac{Na_{ext}}{Km_{ext} + Na_{ext}}\right)\left(\beta(1 - F_{Tcyt})\left(\frac{Km_{cyt}}{Km_{cyt} + Na_{cyt}}\right)\right)\right)$$

$$J_{Tce} = J_{max}\left((F_{Tcyt})\left(\frac{Na_{cyt}}{Km_{cyt} + Na_{cyt}}\right)\left(\frac{1}{\beta}\right)(1 - F_{Text})\left(\frac{Km_{ext}}{Km_{ext} + Na_{ext}}\right)\right)$$

4) **Complex symporter 1: SLC1A1, SLC1A2, SLC1A3, SLC1A6, SLC1A7**

$$J_{Tec} = J_{max}\,(F_{Text})\left(\frac{(Na_{ext})^3}{(Km_{ext})^3 + (Na_{ext})^3}\right)\left(\frac{(H^+{}_{ext})}{Km_{ext} + (H_{ext})}\right)\left(\frac{(K_{cyt})}{Km_{cyt} + (K_{cyt})}\right) * \beta(1 - F_{Tcyt})\left(\frac{(Km_{cyt})^3}{(Km_{cyt})^3 + (Na_{cyt})^3}\right)\left(\frac{Km_{cyt}}{Km_{cyt} + H_{cyt}}\right)\left(\frac{Km_{ext}}{Km_{ext} + K_{ext}}\right)$$

$$J_{Tce} = J_{max}\,(F_{Tcyt})\left(\frac{(Na_{cyt})^3}{(Km_{cyt})^3 + (Na_{cyt})^3}\right)\left(\frac{(H^+{}_{cyt})}{Km_{cyt} + (H_{cyt})}\right)\left(\frac{(K_{ext})}{Km_{ext} + (K_{ext})}\right) * \frac{1}{\beta}(1 - F_{Text})\left(\frac{(Km_{ext})^3}{(Km_{ext})^3 + (Na_{ext})^3}\right)\left(\frac{Km_{ext}}{Km_{ext} + H_{ext}}\right)\left(\frac{Km_{cyt}}{Km_{cyt} + K_{cyt}}\right)$$

5) **Complex symporter 2: SLC6A7, SLC6A9, SLC6A14, SLC6A17, SLC6A18, SLC6A20**

$$J_{Tec} = J_{max}\,(F_{Text})\left(\frac{(Na_{ext})^2}{(Km_{ext})^2 + (Na_{ext})^2}\right)\left(\frac{(Cl^-{}_{ext})}{Km_{ext} + Cl^-_{ext})}\right) * \beta(1 - F_{Tcyt})\left(\frac{(Km_{cyt})^2}{(Km_{cyt})^2 + (Na_{cyt})^2}\right)\left(\frac{Km_{cyt}}{Km_{cyt} + Cl_{cyt}}\right)$$

$$J_{Tce} = J_{max}\,(F_{Tcyt})\left(\frac{(Na_{cyt})^2}{(Km_{cyt})^2 + (Na_{cyt})^2}\right)\left(\frac{(Cl_{cyt})}{Km_{cyt} + (Cl_{cyt})}\right) * \frac{1}{\beta}(1 - F_{Text})\left(\frac{(Km_{ext})^2}{(Km_{ext})^2 + (Na_{ext})^2}\right)\left(\frac{Km_{ext}}{Km_{ext} + Cl_{cyt}}\right)$$

**6) Complex symporter 3: SLC6A5**

$$J_{Tec} = J_{max}\left(F_{Tcyt}\right)\left(\frac{(Na_{ext})^3}{(Km_{ext})^3 + (Na_{ext})^3}\right)\left(\frac{(Cl^{-}{}_{ext})}{Km_{appext} + Cl^{-}_{ext})}\right) * \beta\left(1 - F_{Tcyt}\right)\left(\frac{(Km_{cyt})^3}{(Km_{cyt})^3 + \left(Na_{cyt}\right)^3}\right)\left(\frac{Km_{cyt}}{Km_{cyt} + Cl_{cyt}}\right)\right)$$

$$J_{Tce} = J_{max}\left(F_{Tcyt}\right)\left(\frac{(Na_{cyt})^3}{(Km_{cyt})^3 + (Na_{cyt})^3}\right)\left(\frac{(Cl_{cyt})}{Km_{cyt} + (Cl_{cyt})}\right) * \frac{1}{\beta}\left(1 - F_{Text}\right)\left(\frac{(Km_{ext})^3}{(Km_{ext})^3 + (Na_{ext})^3}\right)\left(\frac{Km_{ext}}{Km_{ext} + Cl_{cyt}}\right)$$

**7) Electroneutral symporter: SLC38A3, SLC38A5**

$$J_{Tec} = J_{max}\left(F_{Text}\right)\left(\frac{(Na_{ext})}{Km_{ext} + Na_{ext}}\right)\left(\frac{\left(H^{+}{}_{cyt}\right)}{Km_{cyt} + (H^{+}_{cyt})}\right)$$

$$J_{Tce} = J_{max}\left(F_{Tcyt}\right)\left(\frac{\left(Na_{cyt}\right)}{Km_{cyt} + (Na_{cyt})}\right)\left(\frac{(H^{+}{}_{ext})}{Km_{ext} + (H^{+}_{ext})}\right)$$

**8) Proton-amino acid symporter: SLC36A1, SLC36A2, SLC36A3, SLC36A4**

$$J_{Tec} = J_{max}\left(F_{Text}\right)\left(\frac{H_{ext}}{Km_{ext} + H_{ext}}\right)\beta\left(1 - F_{Tcyt}\right)\left(\frac{Km_{cyt}}{Km_{cyt} + H_{cyt}}\right)$$

$$J_{Tce} = J_{max}\left(F_{Tcyt}\right)\left(\frac{H_{cyt}}{Km_{cyt} + H_{cyt}}\right)\left(\frac{1}{\beta}\right)\left(1 - F_{Text}\right)\left(\frac{Km_{ext}}{Km_{ext} + H_{ext}}\right)$$

**9) Antiporter: SLC1A4, SLC1A5, SLC7A5, SLC7A8, SLC7A10, SLC7A11, SLC7A12**

$$J_{Tec} = J_{max}(F_{Text})(F_{Tcyt})$$

$$J_{Tce} = J_{Tec}$$

**10) Complex antiporter: SLC7A6, SLC7A7**

$$J_{Tec} = J_{max}(F_{Text})(F_{Tcyt})$$

$$J_{Tce} = J_{max}(F_{Text})(F_{Tcyt})$$

If AAi is cationic: $Fractional\ saturation\ (F_{iext}) = \left(\frac{AAi_{ext}}{Km_{appext}+AAi_{ext}}\right)$ and $F_{icyt} = \left(\frac{AAi_{cyt}}{Km_{appcyt}+AAi_{cyt}}\right)$

IfAAi is neutral: $Fractional\ saturation\ (F_{iext}) = \left(\frac{AAi_{ext}}{Km_{appext}+AAi_{ext}}\right)\left(\frac{Na_{ext}}{Km_{ext}+Na_{ext}}\right)$ and $F_{icyt} = \left(\frac{AAi_{cyt}}{Km_{appcyt}+AAi_{cyt}}\right)\left(\frac{Na_{cyt}}{Km_{cyt}+Na_{cyt}}\right)$

**11) Electrogenic antiporter: SLC7A9**

| | Substrate extracellular | Substrate cytosolic |
|---|---|---|

| | | |
|---|---|---|
| Mode 1 | Cationic | Cationic |
| Mode 2 | Cationic | Neutral |
| Mode 3 | Neutral | Cationic |
| Mode 4 | Neutral | Neutral |

Equation for Mode 1 and 4: $J_{iec} = J_{max}(F_{i1ext})(F_{i2cyt})$

Equation for Mode2: $J_{iec} = J_{max}(F_{i1ext})(F_{i2cyt})\beta$

Equation for Mode 3: $J_{iec} = J_{max}(F_{i1ext})(F_{i2cyt})(\frac{1}{\beta})$

In each case: $J_{aa1ce} = J_{aa2ec}$

### Table S5: Calculated Vmax values for cell lines used in this study

Vmax values are given as nmol/min*mg protein. Red fields were excluded from averages.

| Transporter | Probe AA | | | A549 | | |
|---|---|---|---|---|---|---|
| SLC1 | | Conc | min | Activity | Km | Vmax |
| EAAT3 | Glu | 100 | 6 | | 62 | |
| EAAT1 | Glu | 100 | 6 | | 48 | |
| EAAT2 | Glu | 100 | 6 | | 48 | |
| EAAT4 | Glu | 100 | 6 | | 3 | |
| EAAT5 | Glu | 100 | 6 | | 64 | |
| ASCT1 | Ala | 300 | 2 | | 71 | |
| | Pro | 100 | 2 | 1.2 | 672 | 4.632 |
| ASCT2 | Ala | 300 | 2 | 48 | 18 | 25.44 |
| | Gln | 100 | 2 | 23 | 24 | 14.26 |
| | Gly | 100 | 6 | 1.5 | 361 | 1.1525 |
| SLC6 | | | | | | |
| GlyT2 | Gly | 100 | 6 | | 17 | |
| GlyT1 | Gly | 100 | 6 | | 220 | |
| PROT | Pro | 100 | 6 | | 6 | |
| ATB0,+ | Leu | 100 | 2 | | 12 | |
| | Arg | 100 | 6 | | 104 | |
| B0AT2 | Leu | 100 | 2 | | 81 | |
| | Ala | 300 | 2 | | 670 | |
| | Pro | 100 | 6 | | 195 | |
| NTT4 | Gln | 100 | 2 | | 5200 | |
| | Leu | 100 | 2 | | 280 | |
| | Pro | 100 | 6 | | 360 | |
| B0AT1 | Leu | 100 | 2 | | 1000 | |
| | Ala | 300 | 2 | | 4100 | |
| | Gln | 100 | 2 | | 3200 | |
| SIT1 | Pro | 100 | 6 | | 130 | |
| SLC7 | | | | | | |
| CAT1 | Arg | 100 | 6 | 10 | 77 | 2.95 |
| CAT2 | Arg | 100 | 6 | | 2700 | |
| CAT3 | Arg | 100 | 6 | | 103 | |
| LAT1 | Leu | 100 | 2 | 26 | 20 | 15.6 |
| | Gln | 100 | 2 | 3.2 | 1640 | 27.84 |
| y+LAT2 | Leu | 100 | 2 | 6.4 | 236 | 10.752 |
| | Arg | 100 | 6 | 15 | 138 | 5.95 |
| y+LAT1 | Leu | 100 | 2 | | 44 | |
| | Arg | 100 | 6 | | 340 | |
| LAT2 | Leu | 100 | 2 | 12 | 119 | 13.14 |
| | Ala | 300 | 2 | 11.3 | 187 | 9.171833 |
| | Gln | 100 | 2 | 6.8 | 151 | 8.534 |
| b0,+AT | Leu | 100 | 2 | | 107 | |
| | Arg | 100 | 6 | | 108 | |
| Asc-1 | Ala | 300 | 2 | | 23 | |
| xCT | Glu | 25 | 6 | 4.8 | 92 | 3.744 |
| AGT1 | Glu | 100 | 6 | | 21 | |
| SLC16 | | | | | | |
| TAT1 | Phe | 100 | | | 7020 | |
| SLC36 | | | | | | |
| PAT1 | Gly | 100 | 6 | | 7000 | |
| | Pro | 100 | 6 | | 2800 | |
| PAT2 | Gly | 100 | 6 | | 590 | |
| | Pro | 100 | 6 | | 120 | |
| PAT4 | Pro | 100 | 6 | | 3 | |
| | Ala | 300 | 2 | | 1480 | |
| SLC38 | | | | | | |
| SNAT1 | Ala | 300 | 2 | 9 | 306 | 9.09 |
| | Gln | 100 | 2 | 4.5 | 230 | 7.425 |
| | Pro | 100 | 6 | 7.8 | 1000 | 14.3 |
| | Gly | 100 | 6 | 2.1 | 890 | 3.465 |
| SNAT2 | Ala | 300 | 2 | 1.3 | 529 | 1.796167 |
| | Gln | 100 | 2 | 0.6 | 1650 | 5.25 |
| | Leu | 100 | 2 | 5.3 | 1000 | 29.15 |
| | Pro | 100 | 6 | 3.6 | 800 | 5.4 |
| | Gly | 100 | 6 | 0.2 | 800 | 0.3 |
| SNAT3 | Gln | 100 | 2 | | 1500 | |
| SNAT4 | Ala | 300 | 2 | | 4200 | |
| | Pro | 100 | 6 | | | |
| | Gln | 100 | 2 | | 2500 | |
| | Arg | 100 | 6 | 3.6 | 300 | 2.4 |
| SNAT5 | Ala | 300 | 2 | 6.9 | 2500 | 32.2 |
| | Gln | 100 | 2 | 4.3 | 3200 | 70.95 |
| | Gly | 100 | 6 | | 15200 | |
| SNAT7 | Gln | 100 | 2 | | | |
| SNAT8 | Gln | 100 | 2 | | | |
| SLC43 | | | | | | |
| LAT3 | Leu | 100 | 2 | | 1024 | 0 |
| | Phe | 100 | 2 | | 1206 | |
| LAT4 | Leu | 100 | 2 | | 3733 | |
| | Phe | 100 | 2 | | 4694 | |

| Transporter | Probe AA | | | U87-MG | | |
|---|---|---|---|---|---|---|
| SLC1 | | Conc | min | Activity | Km | Vmax |
| EAAT3 | Glu | 100 | 6 | 2.5 | 62 | 0.675 |
| EAAT1 | Glu | 100 | 6 | | 48 | |
| EAAT2 | Glu | 100 | 6 | | 48 | |
| EAAT4 | Glu | 100 | 6 | | 3 | |
| EAAT5 | Glu | 100 | 6 | | 64 | |
| ASCT1 | Ala | 300 | 2 | | 71 | |
| | Pro | 100 | 2 | 0.9 | 672 | 3.474 |
| ASCT2 | Ala | 300 | 2 | 43.4 | 18 | 23 |
| | Gln | 100 | 2 | 24.7 | 24 | 15.31 |
| | | | | | 361 | |
| SLC6 | | | | | | |
| GlyT2 | Gly | 100 | 6 | | 17 | |
| GlyT1 | Gly | 100 | 6 | | 220 | |
| PROT | Pro | 100 | 2 | | 6 | |
| ATB0,+ | Leu | 100 | 2 | | 12 | |
| | Arg | 100 | 6 | | 104 | |
| B0AT2 | Leu | 100 | 2 | | 81 | |
| | Ala | 300 | 2 | | 670 | |
| | Pro | 100 | 2 | | 195 | |
| NTT4 | Gln | 100 | 2 | | 5200 | |
| | Leu | 100 | 2 | | 280 | |
| | Pro | 100 | 2 | | 360 | |
| B0AT1 | Leu | 100 | 2 | | 1000 | |
| | Ala | 300 | 2 | | 4100 | |
| | Gln | 100 | 2 | | 3200 | |
| SIT1 | Pro | 100 | 2 | | 130 | |
| SLC7 | | | | | | |
| CAT1 | Arg | 100 | 6 | | 77 | |
| CAT2 | Arg | 100 | 6 | | 2700 | |
| CAT3 | Arg | 100 | 6 | | 103 | |
| LAT1 | Leu | 100 | 2 | 26 | 20 | 15.6 |
| | Gln | 100 | 2 | | 1640 | |
| y+LAT2 | Leu | 100 | 2 | 6.4 | 236 | 55.68 |
| | Arg | 100 | 6 | 4.3 | 138 | 2.408 |
| y+LAT1 | Leu | 100 | 2 | | 44 | |
| | Arg | 100 | 6 | | 340 | |
| LAT2 | Leu | 100 | 2 | 12 | 119 | 26.4 |
| | Ala | 300 | 2 | 4.8 | 187 | 3.352 |
| | Gln | 100 | 2 | 3.8 | 151 | 5.453 |
| b0,+AT | Leu | 100 | 2 | | 107 | |
| | Arg | 100 | 6 | | 108 | |
| Asc-1 | Ala | 300 | 2 | | 23 | |
| xCT | Glu | 100 | 6 | 10 | 92 | 2.05 |
| AGT1 | Glu | 100 | 6 | | 21 | |
| SLC16 | | | | | | |
| TAT1 | Phe | 100 | | | 7020 | |
| SLC36 | | | | | | |
| PAT1 | Gly | 100 | 6 | 0.5 | 7000 | 0.083 |
| | Pro | 100 | 6 | | 2800 | |
| PAT2 | Gly | 100 | 6 | | 590 | |
| | Pro | 100 | 6 | | 120 | |
| PAT4 | Pro | 100 | 6 | 0.6 | 3 | 0.22 |
| | Ala | 300 | 2 | | 1480 | |
| SLC38 | | | | | | |
| SNAT1 | Ala | 300 | 2 | 7.4 | 306 | 21.95 |
| | Gln | 100 | 2 | | 230 | |
| | Pro | 100 | 6 | | 1000 | |
| | Gly | 100 | 6 | | 890 | |
| SNAT2 | Ala | 300 | 2 | 4 | 529 | 4.04 |
| | Gln | 100 | 2 | 3.6 | 1650 | 5.94 |
| | Leu | 100 | 2 | 5.3 | 1000 | 29.15 |
| | Pro | 100 | 6 | | 800 | |
| | Gly | 100 | 6 | | 800 | |
| SNAT3 | Gln | 100 | 2 | | 1500 | |
| SNAT4 | Ala | 300 | 2 | | | |
| | Pro | 100 | 6 | | 4200 | |
| | Gln | 100 | 2 | | 2500 | |
| | Arg | 100 | 6 | 2 | 300 | 3.667 |
| SNAT5 | Ala | 300 | 2 | 4.7 | 2500 | 8.617 |
| | Gln | 100 | 2 | 3.3 | 3200 | 14.85 |
| | Gly | 100 | 2 | | 15200 | |
| SNAT7 | Gln | 100 | 2 | | | |
| SNAT8 | Gln | 100 | 2 | | | |
| SLC43 | | | | | | |
| LAT3 | Leu | 100 | 2 | | 1024 | |
| | Phe | | | | 1206 | |
| LAT4 | Leu | 100 | 2 | | 3733 | |
| | Phe | | | | 4694 | |
