## Supplemental figures for "A unified model of amino acid homeostasis in mammalian cells"

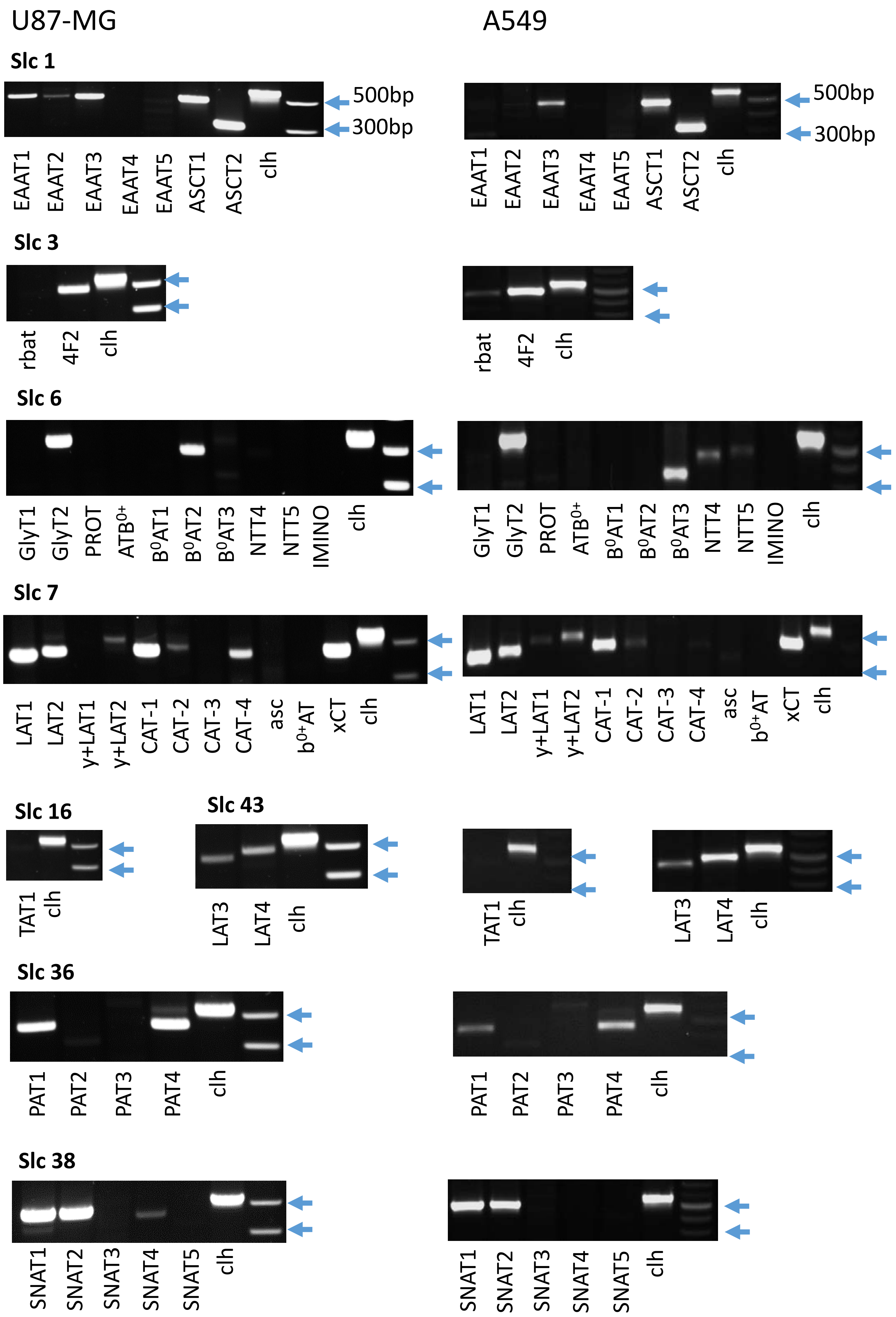

U87-MG
A549
Slc 1
500bp
300bp
EAAT1
EAAT2
EAAT3
EAAT4
EAAT5
ASCT1
ASCT2
clh
Slc 3
rbat
4F2
clh
Slc 6
GlyT1
GlyT2
PROT
ATB$^{0+}$
B$^{0}$AT1
B$^{0}$AT2
B$^{0}$AT3
NTT4
NTT5
IMINO
clh
Slc 7
LAT1
LAT2
y+LAT1
y+LAT2
CAT-1
CAT-2
CAT-3
CAT-4
asc
b$^{0+}$AT
xCT
clh
Slc 16
TAT1
clh
Slc 43
LAT3
LAT4
clh
Slc 36
PAT1
PAT2
PAT3
PAT4
clh
Slc 38
SNAT1
SNAT2
SNAT3
SNAT4
SNAT5
clh

Fig. S1: Analysis of transporter expression by RT-PCR.
For reverse transcription 2 µg of total RNA isolated from U87-MG cells or A549 cells was used in a reaction volume of 20 uL. One µl of the cDNA sample was used for a 30 cycle PCR reaction. Predicted fragment sizes are listed in the methods section.

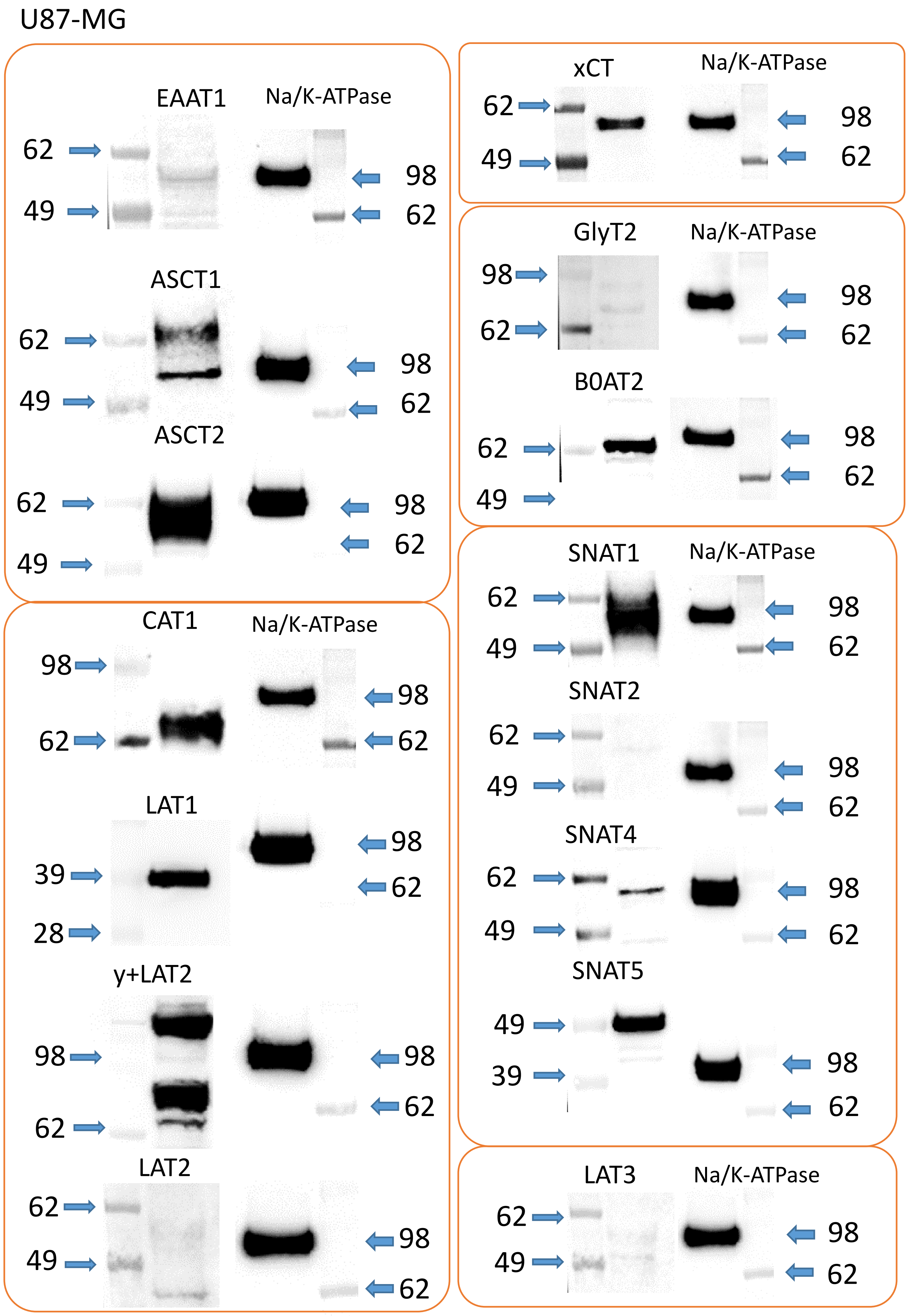
U87-MG
EAAT1
Na/K-ATPase
62
49
98
62
ASCT1
62
49
98
62
ASCT2
62
49
98
62
CAT1
Na/K-ATPase
98
62
98
62
LAT1
39
28
98
62
y+LAT2
98
62
98
62
LAT2
62
49
98
62
xCT
Na/K-ATPase
62
49
98
62
GlyT2
Na/K-ATPase
98
62
98
62
B0AT2
62
49
98
62
SNAT1
Na/K-ATPase
62
49
98
62
SNAT2
62
49
98
62
SNAT4
62
49
98
62
SNAT5
49
39
98
62
LAT3
Na/K-ATPase
62
49
98
62

Fig S2. Surface expression of amino acid transporters in U87-MG cells.
Surface biotinylation was performed before detection of amino acid transporters by western blotting. Antibodies were used as listed in table M5 (Methods & Materials). For comparison $Na^+/K^+$-ATPase was detected on the same blot after stripping.

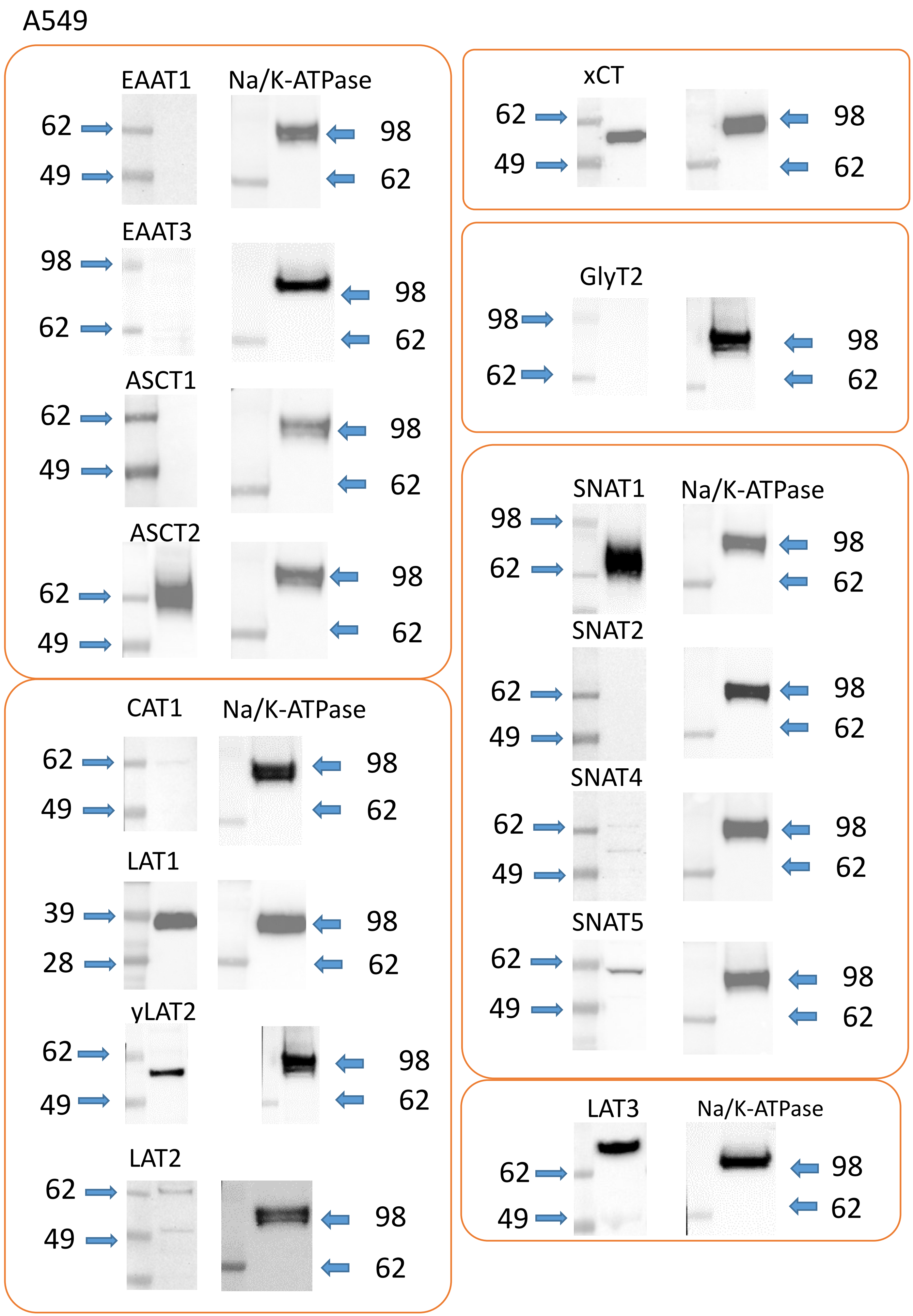
A549
EAAT1
Na/K-ATPase
62
49
98
62
EAAT3
98
62
98
62
ASCT1
62
49
98
62
ASCT2
62
49
98
62
CAT1
Na/K-ATPase
62
49
98
62
LAT1
39
28
98
62
yLAT2
62
49
98
62
LAT2
62
49
98
62
xCT
62
49
98
62
GlyT2
98
62
98
62
SNAT1
Na/K-ATPase
98
62
98
62
SNAT2
62
49
98
62
SNAT4
62
49
98
62
SNAT5
62
49
98
62
LAT3
Na/K-ATPase
62
49
98
62

Fig S3. Surface expression of amino acid transporters in A549 cells.
Surface biotinylation was performed before detection of amino acid transporters by western blotting. Antibodies were used as listed in table M5 (Methods & Materials). For comparison $Na^{+}/K^{+}$-ATPase was detected on the same blot after stripping.

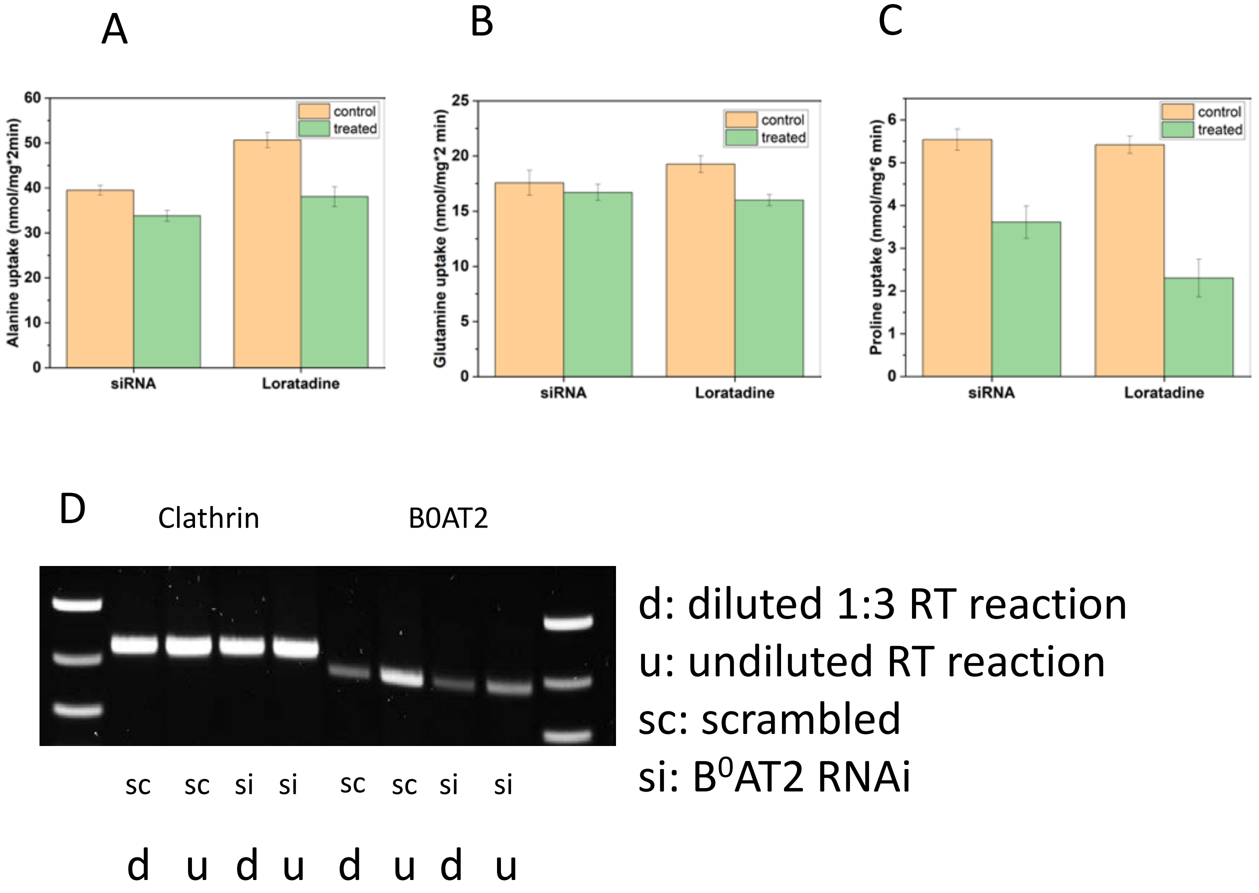


Fig. S4: Contribution by $B^0AT2$ to amino acid transport in U87-MG cells. Alanine (A), glutamine (B) and proline (C) transport was measured in control cells and in cells where $B^0AT2$ was silenced by RNAi. For comparison, inhibition by the published $B^0AT2$ inhibitor loratadine is shown. Silencing of $B^0AT2$ mRNA is shown in (D), using 1:3 diluted and undiluted reverse transcriptase reactions for PCR.

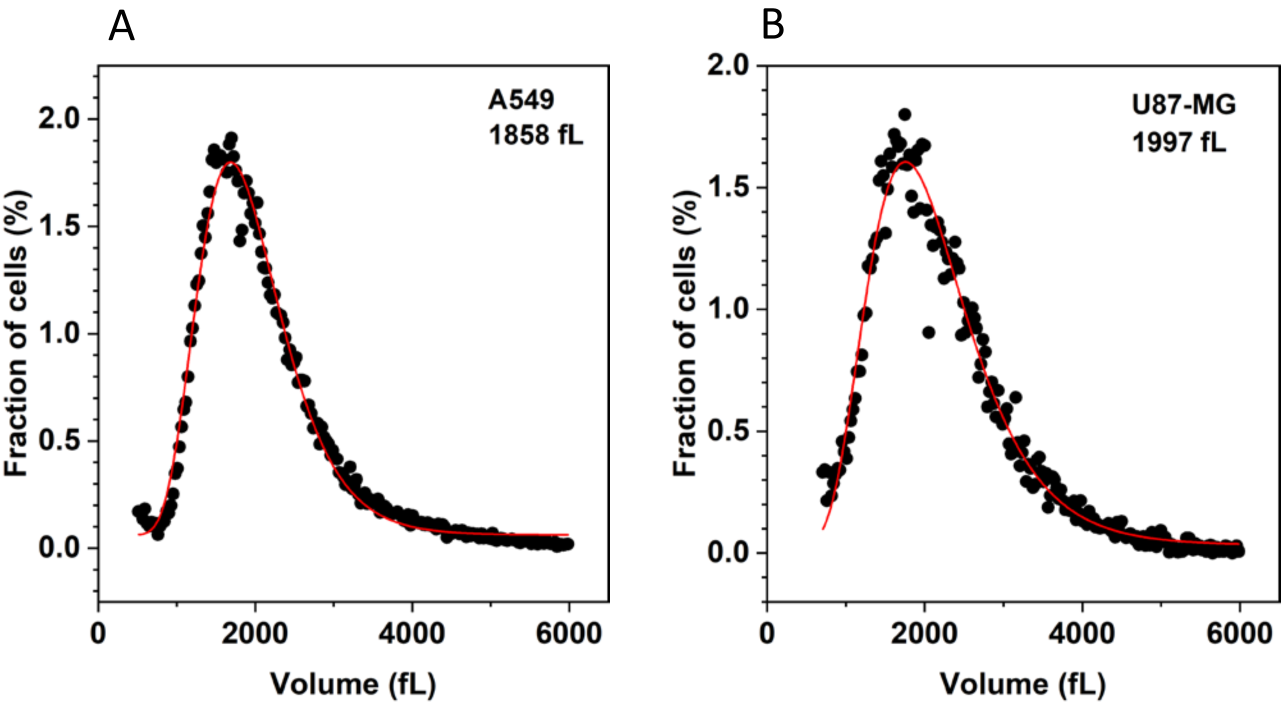


Fig. S5: Volume analysis of cell lines.
Cell lines A549 (A) and U87-MG (B) were dislodged by trypsin treatment and dispersed by passage through 70 µm sieve. Cell size was measured by a Multisizer 4 Coulter counter.

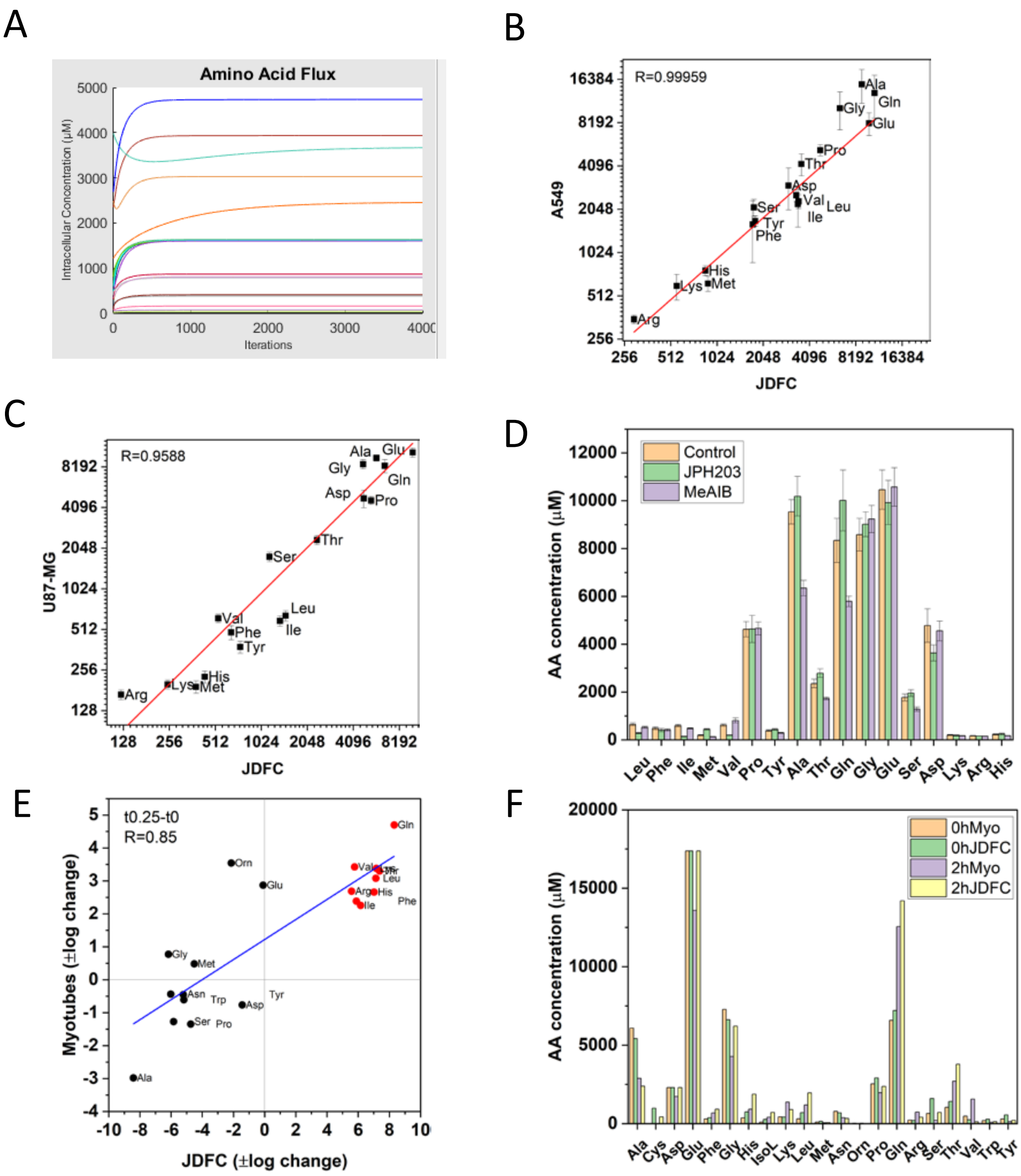


Fig. S6: Modelling of amino acid transport

A) Snapshot of JDFC demonstrating a stable equilibrium of cytosolic amino acid concentrations starting from initial concentrations. B,C) Correlation between experimental and simulated amino acid concentrations in A549 cells (B) and U87-MG cells (C). D) Effect of amino acid transport inhibitors JPH203 and MeAIB on cytosolic amino acid concentrations in U87-MG cells after 1h incubation. E) Correlation between experimental and simulated dynamic changes (log scale) in myotubes after incubation with an amino acid mixture similar to AXA2678 for 15 min. Supplemented amino acids are indicated by red dots. F) Experimental and simulated cytosolic amino acid concentrations in human myotubes grown in sarcopenia-inducing medium after incubation with AXA2678 for 2h. Sample amino acid concentrations were multiplied x100 to be in the range of the simulated data.
